# CXCL17 binds efficaciously to glycosaminoglycans with the potential to modulate chemokine signalling

**DOI:** 10.1101/2023.07.07.548106

**Authors:** Sean P. Giblin, Sashini Ranawana, Shyreen Hassibi, Holly L. Birchenough, Kyle T. Mincham, Robert J. Snelgrove, Tomoko Tsuchiya, Shiro Kanegasaki, Douglas Dyer, James E. Pease

## Abstract

CXCL17 is a mucosally secreted protein, and the most recently identified human chemokine, an assignment based on protein fold prediction and chemotactic activity for leukocytes. However, these credentials have been the subject of much recent discussion and no experimental evidence has been presented regarding the definitive structure of CXCL17. In this study, we evaluated the structural and chemoattractant credentials of CXCL17 to better characterise this molecule, and gain deeper insights into its functional role as a glycosaminoglycan (GAG) binding protein.

In the absence of structural information, *in silico* modelling techniques assessed the likelihood of CXCL17 adopting a chemokine-fold. Recombinant CXCL17 was synthesized in mammalian and prokaryotic systems. Modified Boyden chamber and real-time chemotaxis assays assessed the ability of CXCL17 to promote chemotaxis of murine splenocytes, human neutrophils and CXCR1-transfectants. The efficacy of CXCL17 binding to GAGs was quantified with solid-phase assays and bio-layer interferometry techniques.

All modelling efforts failed to support classification of CXCL17 as a chemokine based on its predicted conformation. Recombinant CXCL17 was observed to dimerize as a function of concentration, a characteristic of several chemokines. Contrary to a previous report, CXCL17 was not chemotactic for murine splenocytes, although it was a low-potency chemoattractant for human neutrophils at micromolar concentrations, several orders of magnitude higher than those required for CXCL8. As anticipated due to its highly basic nature, CXCL17 bound to GAGs robustly, with key C-terminal motifs implicated in this process. While inactive via CXCR1, CXCL17 was found to inhibit CXCR1-mediated chemotaxis of transfectants to CXCL8 in a dose-dependent manner.

In summary, despite finding little evidence for chemokine-like structure and function, CXCL17 readily bound GAGs, and could modulate chemotactic responses to another chemokine *in vitro.* We postulate that such modulation is a consequence of superior GAG-binding, and that C-terminal fragments of CXCL17 may serve as prototypic inhibitors of chemokine function.

## 1 Introduction

Chemokines are a family of around 40 small soluble cytokines in humans, noted for their chemoattractant properties, particularly for leukocytes [1]. Despite quite varied degrees of homology, chemokines adopt the same characteristic ‘Greek-key’ protein fold, consisting of three anti-parallel β-sheets overlaid by a C-terminal α-helix [2]. The vast majority of chemokine family members fall into two classes known as CC and CXC chemokines, which describes the arrangement of conserved N-terminal cysteine residues which are either adjacent or interspersed with a single amino acid [3]. As is the case with other leukocyte chemoattractants such as C5a and fMLP, chemokines exert their effects by binding to members of the G protein-coupled receptor (GPCR) superfamily expressed on the leukocyte surface [4; 5; 6].

CXCL17 was originally identified as a potential chemokine on the basis of apparent homology with CXCL8 as a result of molecular modelling efforts [7]. However, the validity of this claim has recently been questioned by ourselves and others [8; 9]. In a notable departure from most other chemokines, the primary structure of CXCL17 contains six conserved cysteine residues, a feature seen in only one other CXC chemokine, CXCL16 [10], and in the 6-Cys subset of CC chemokines comprised of CCL1 [11], CCL15 [12], CCL21 [13], CCL23 [14] and CCL28 [15]. CXCL17 is constitutively expressed at mucosal surfaces such as those in the proximal digestive tract, lung, and stomach [16; 17], although its precise function at these locations is not fully understood.

The mature CXCL17 protein sequence contains many arginine and lysine residues and as a result is predicted to be highly positively charged at a physiological pH with an isoelectric point (pI) of 10.95. This ranks it among the most cationic of CXC chemokines [18]. This highly basic nature suggests CXCL17 may play a role as an antimicrobial peptide, and *in vitro* data have been reported showing a broad spectrum of microbicidal activity against bacteria and fungi [16]. Previous studies reported CXCL17 to be a chemoattractant for monocytes and dendritic cells [7; 17] and murine splenocytes [19]. Deletion of the *cxcl17* gene in mice was observed to result in reduced numbers of alveolar macrophages within the lungs, which has led to suggestions of a role for CXCL17 in their recruitment [20]. The GPCR known as GPR35 is expressed by some subsets of leukocytes and was postulated to be a receptor for CXCL17 [21], although this has been challenged by ourselves and others [22; 23]. Notably, GPR35 was recently de-orphanized and identified as a receptor for the chemotactic serotonin metabolite, 5-hydroxyindoleacetic acid and shown to recruit GPR35^+^ neutrophils at low nanomolar concentrations [24].

A currently unexplored possibility is that the high isoelectric point for CXCL17 may confer an ability to interact with extracellular matrix proteoglycans and glycosaminoglycans (GAGs) in the mucosae. GAGs are comprised of heterogenous sub-populations of linear, highly sulfated polysaccharides which via covalent interactions with proteoglycans reside in close association to cell membranes [25; 26]. Chemokine-GAG interactions are widely reported elsewhere and are essential for retaining chemokine gradients on specific tissues and cell surfaces, while promoting and decoding chemokine signals that drive leukocyte motility, survival and function (reviewed in [27; 28]). However, the potential for GAG interactions with CXCL17 and their functional consequences are not yet known.

In this study, we report the chemotactic activity of CXCL17 for human neutrophils assessed using a real-time assay method, and the likelihood of CXCL17 adopting a chemokine-like fold using recent advances in molecular modelling. We also highlight the ability of CXCL17 to bind to GAGs and the effect of CXCL17 on the chemotactic responses of CXCR1 transfectants to CXCL8. Based upon these findings, we suggest that CXCL17 may have modest chemotactic activity for neutrophils and that it can modulate the activity of other chemokines which depend upon glycosaminoglycan binding for their activity.

## 2 Materials and Methods

### 2.1 Materials

Materials were purchased from SIGMA-Aldrich (Poole, UK) unless otherwise stated. Oligonucleotide production and DNA sequencing services were from MWG-Biotech (Ebersberg, Germany). HPLC materials were from Cytiva (Amersham, UK). Recombinant proteins were from R&D systems (Bio-Techne Ltd., Abingdon, UK). Heparin dp8 and heparan sulfate used in the bio-layer interferometry (BLI) assays were from Iduron (Alderley Edge, UK). Primary antibodies used in western blotting were sheep anti-human CXCL17 pAb (#AF4207), mouse anti-CXCL17 mAb (#MAB4207), mouse anti-human CXCL4 mAb (#MAB7951) all from Bio-Techne. These were detected with either protein G-conjugated HRP or goat anti-mouse IgG Alexa Fluor Plus 800n, both from Thermofisher, (Paisley, UK).

### 2.2 Signal peptide prediction, phylogenetic analysis and prediction of glycosylation sites

Signal peptide cleavage prediction was performed on the full-length amino acid sequence for human CXCL17 (1-119) (Protein IUD Q6UXB2), using the Signal-P 6.0 server [29]. CXCL17 was aligned against all other human CXC chemokines using SnapGene® software (Dotmatic, available at <u>snapgene.com</u>). Phylogenetic analysis was performed by EMBL-EBI MUSCLE alignment [30], with the results extracted and displayed as a midpoint rooted neighbour joining tree without distance corrections using interactive Tree Of Life [31]. Secondary structure predictions were performed with the DSC server [32], showing predictions for CXCL8 (28-99) (P10145) and CXCL17 (24-119). Putative *N-*linked and *O-*linked glycosylation sites were predicted using the NetNGlyc-1.0 [33] and NetOGlyc-4.0 [34] servers respectively.

### 2.3 *In silico* structural modelling of CXCL17

The CXCL17 (24-119) tertiary structure was predicted by AlphaFold2 (DeepMind, EMBL-EBI) using the ColabFold interface as described in [35]. Target sequence was uploaded as a pdb70 template, and multiple sequence alignment (MSA) was performed by MMSeqs2 against Uniref and Environmental structure libraries with unpaired and paired sequences. Structural modelling was performed with AlphaFold2-ptm and AlphaFold multimer v2 predictions, with 3-, 12-, and 48-iterances of model recycling. A CXCL17 (24-119) homodimer structure was modelled using the same process with 12-iterances of model recycling. Stereochemical plausibility and confidence in the model expressed in the predicted local distance test (plDDT) scores per residue [36], and by the Predicted Aligned Error (PAE) measured in Å distance [37]. *De novo* folding of CXCL17 (24-119) was performed by specifying no template mode and single sequence MSA mode for 48 model recycles. Predictions of the CXCL17 (24-119) structure by RoseTTAFold with ColabFold was performed by MMSeqs2 as described [35], and *de novo* folding was performed using single sequence input. Predictions were run by RoseTTAFold for mainchain, and Scrwl4 for sidechain predictions. Structural predictions generated by C-I-TASSER integrated the I-TASSER hierarchical structure modelling approach with deep learning based contact predictions to guide Replica Exchange Monte Carlo (REMC) simulations to produce CXCL17 structure models as described [38].

### 2.4 Generation of CXCL4, CXCL17 and SUMO3-CXCL17 mutant constructs

The pE-SUMOpro3 AMP vector (Lifesensors Inc, PA, USA) was modified by site-directed mutagenesis to insert a silent *AgeI* restriction site in the final codon of the SUMO3 open reading frame (ORF). This facilitated subsequence cloning of chemokine inserts. The vector was renamed in house as pEM-SUMOpro3 AMP. Open reading frames (ORFs) encoding the CXCL4 and CXCL17 (24-119) sequences without predicted signal peptides were subcloned into the pEM-SUMOpro3 AMP vector at the *AgeI* and *BamHI sites.* A panel of SUMO3-CXCL17 truncation mutants was generated by SDM by introducing stop codons at the required positions using the QuikChange II Site-Directed Mutagenesis Kit (Agilent Technologies; Santa Clara, USA).

### 2.5 Expression and purification of recombinant proteins

SUMO3-CXCL4 was expressed as inclusion bodies in BL21(DE3) pLysS *E.*coli, as previously described [39]. WT SUMO3-CXCL17 and variants were expressed as soluble proteins in SHuffle® T7 Competent *E.coli* (New England Biolabs, Hitchin, UK). In both protocols, protein was induced by the addition of 100 mM Isopropyl ß-D-1-thiogalactopyranoside (IPTG) when cultures had reached log phase. Cultures were grown for a further 5 hours after which pellets were harvested. Pellets were either stored at -20°C until further use or immediately lysed by sonication in HisTrap buffer A (50 mM Tris pH 8.0, 150 mM NaCl, 10% glycerol, 20 mM imidazole) supplemented with cOmplete™ Mini EDTA-free protease inhibitor cocktail. Lysates were clarified by centrifugation at 21,000 x g for 20 minutes followed by further clarification by 0.22 µm filtration.

Purifications were performed using an ÄKTA Pure Protein Purification System (Cytiva). CXCL4 was purified from inclusion bodies following solubilization in chaotropic buffer A (50mM Tris pH 8.0, 6M GuHCl, 50 mM NaCl, 20mM Imidazole) at room temperature (RT) overnight. Solubilised CXCL4 was loaded on a HisTrap HP 5ml FF crude column and washed with 5 column volumes (CV) of chaotropic buffer A, and eluted with 0-100% chaotropic buffer B (50mM Tris pH 8.0, 6M GuHCl, 50 mM NaCl, 500mM Imidazole). The lysates containing SUMO3-CXCL17 and variants were loaded on HisTrap HP 5ml FF crude column pre-equilibrated with HisTrap buffer A. The column was washed with 20 CV of the same buffer, followed by a gradient of 0-100% HisTrap Buffer B (50 mM Tris pH 8.0, 150 mM NaCl, 10% glycerol, 500 mM imidazole). Eluates were pooled and diluted to 0.5 mg/ml.

The SUMO3 tag was removed by digestion with a preparation of *Saccharomyces cerevisiae* Ubiquitin-like-specific protease 1 (*UlpI) which was* expressed in *E. coli* using a pET-28a construct and purified via an N-terminal 6xHis tag as previously described [40]. For the SUMO3-CXCL4 construct, eluate was dialysed into digestion buffer (50mM Tris pH8.0, 150mM NaCl, 10% glycerol, 1mM DTT) at 4°C over 3 days. *UlpI* and SUMO3-CXCL4 were incubated at a 1:30 mass ratio at 4°C, with gentle stirring for 24 hours. For CXCL17, *UlpI* and SUMO3-CXCL17 were incubated at a 1:25 mass ratio, in a non-reducing digestion buffer at 4°C for 24 hours. Removal of the SUMO3-tag resulted in CXCL17 precipitating. The CXCL17 precipitate was subsequently solubilized in chaotropic buffer at RT for 24 hours.

Separation of *UlpI* and the SUMO3 tag from free chemokine was performed on a HisTrap HP 1ml column, pre-equilibrated with chaotropic buffer A supplemented with 1mM DTT, and CXCL17 eluted with chaotropic buffer A. Eluate was dialysed into 50 mM Tris pH 9.0, 150mM NaCl, 10% glycerol, 1mM DTT at 4 °C overnight. Purified CXCL4 and CXCL17 were refolded by ‘infinite’ dilution [41] in cysteine/cystine, refolding buffer overnight at 4 °C. Refolded CXCL4 and CXCL17 were purified on a HiTrap Heparin HP 1ml column, washed with 20CV HiTrap buffer A (50mM Tris pH8.0, 150mM NaCl, 10% glycerol) and eluted with 10CV 0-100% gradient of HiTrap Buffer B (50mM Tris pH8.0, 2.5M NaCl, 10% glycerol). Eluates were dialysed into storage buffer (50mM Tris pH 8.0, 150 mM NaCl, 10% glycerol) and snap frozen in liquid nitrogen for storage. The molecular mass and purity of CXCL4, CXCL17, SUMO3-CXCL17 and mutants was confirmed by SDS-PAGE, and identity confirmed by Western blot. Concentration was assessed by human CXCL4 DuoSet ELISA (DY795), or SDS-PAGE followed by densitometry.

### 2.6 Cell culture and transfection

CHO-761H cells [42] were seeded in 24-well plates, at 2x10^5^ cells/well in 24-Ham’s F12 media, supplemented with glutamine, penicillin/streptomycin (PS) and 10% FBS. After resting for 24 hours, cells were transfected with pmaxGFP (500ng/µl) or pcDNA3.1-CXCL17 (194.1 ng/µl) using lipofectamine 3000 and were cultured in Opti-MEM media containing 1% PS. Mock treated cells underwent the same transfection conditions in the absence of plasmid. After 48 hours, cell supernatants (SN) were collected and cells were lysed in PBS, 1% IGEPAL, 0.4% iodoacetamide, 0.4% EDTA, 2% protease inhibitors. pcDNA3.1-CXCL17 and mock SN was concentrated with 50μL HisPur™ Ni-NTA Resin (ThermoFisher). Transfected cells were analysed by flow cytometry to confirm transfection efficacy and protein expression after 48 hours. Cells were detached with versene, resuspended in FACS buffer containing 1:10000 TO-PRO-3 iodine, and samples were read with a FASCalibur flow cytometer (Becton Dickinson, UK), according to manufacturer’s instructions. To inhibit *O-*linked glycosylation, transfected CHO-761H cells were treated with 1 - 2mM of Benzyl-GalNac (BGN), or DMSO vehicle control one hour after transfection. Lysates were collected after 48 hours and assessed by SDS-PAGE and Western blot.

Primary human neutrophils were isolated from healthy donor whole blood samples obtained with local ethical approval. Purification used the MACSxpress^®^ Whole Blood Neutrophil Isolation Kit (Miltenyi) according to the manufacturer’s instructions. Prior to chemotaxis assessment, neutrophils were resuspended to a concentration of 5x10^6^ cells/mL in chemotaxis buffer RPMI-1640, 1% P/S, 0.1% BSA and were rested for one hour at 37°C. Murine IL-3 dependent pro-B-cells Ba/F3 were maintained in RPMI-1640, 10% FBS, 1% PS, 50µM 2-β-ME, supplemented with 1 ng/ml mouse IL-3 (Peprotech). Stable hCXCR1 expressing Ba/F3 cells were generated by transfection in a 0.4cm-electroporation cuvette with 50ul 10 mg/ml tRNA, with 1µg pcDNA3.1-CXCR1 plasmid at 330V, 950µF using a MicroPulser electroporator (BioRad). Expression of CXCR1 was confirmed by flow cytometry as previously described [43], and maintained by antibiotic resistance selected by 400 µg/ml G418.

### 2.7 Isolation of mouse splenocytes

Eight-to 10-week old female C57BL/6 mice were purchased from Charles River (Oxford, UK) and housed at Imperial College Central Biomedical Services facility. Mice were kept in specific pathogen-free conditions and provided autoclaved food, water and bedding ad libitum. All animal procedures were performed in accordance with the recommendations in the Guide for the Use of Laboratory Animals of Imperial College London. All animal procedures and care conformed strictly to the UK Home Office Guidelines under the Animals (Scientific Procedures) Act 1986, and the protocols were approved by the Home Office of Great Britain.

### 2.8 Chemotaxis assays

Real-time chemotaxis assessment with the TAXIScan-12 system 2 (Hirata Corp., Japan) was performed as previously described [44]. 5x10^2^ neutrophils were loaded onto the instrument in each channel, aligned and then allowed to migrate for 60 minutes along chemokine gradients generated by the addition of 1µl of recombinant 10 nM CXCL8 or varying concentrations of CXCL17 (24-119) purchased from R&D Systems. Basal migration was recorded in the absence of stimulus. Images were captured every 60 seconds and individual migration paths were manually tracked using the manual tracking function of ImageJ1.46. Chemotaxis was quantified using the IBIDI chemotaxis tool [45] and was expressed as directionality, velocity (µm/min), and y-axis directional forward migration index (yFMI). yFMI describes the forward migration of cells parallel to the chemokine gradient and is calculated by dividing the Δy value of a cell track end-point, by the total accumulated distance travelled.

Modified Boyden chamber assays were performed as previously described [23] using 96-well 5 µm pore CHEMO Tx^®^ plates (Neuro Probe; Gaithersburg, MD) [43]. Dose responses to murine CXCL17 alone or to 10% FBS were performed with C57BL/6 murine splenocytes. Dose responses to human CXCL8 and CXCL17 alone or in combination were generated using Ba/F3 cells stably expressing CXCR1. Cells were suspended in chemotaxis buffer and migrated for 5 hours at 37°C, 5% CO_2_ in a humidified chamber after which cells traversing the membrane were assessed by CellTiterGlo^®^ Luminescent Cell Viability Assay (Promega; Madison, WI) according to the manufacturers protocol using a TopCount^®^ NXT^TM^ Microplate Scintillation and Luminescence Counter. Data were normalized to the total luminescence from an input of 2x10^5^ cells. Chemotactic indices were calculated by dividing the luminescence of cells migrating to stimulus, by the basal level of migration.

### 2.9 SDS-PAGE and Western blot

Proteins were resolved by reducing SDS-PAGE, using 12% NuPAGE Bis-Tris Mini Gels and MES buffer on a Mini Gel Tank (Invitrogen) according to manufacturer’s instructions. Proteins were transferred to nitrocellulose iBlot™ mini transfer stacks, with an iBlot™ gel transfer device and membranes were blocked with PBS containing 0.05% Tween-20 (PBS-T) and 5% (w/v) milk powder for 1 hour and probed with 0.1 µg/ml of the relevant primary antibody in fresh blocking buffer overnight at 4 °C with agitation. Blots were washed 3 times in PBS-T and probed with 0.2µg/ml protein G-conjugated HRP or 0.1µg/ml diluted goat anti-mouse IgG Alexa Fluor Plus 800 in blocking buffer at RT for 2 hours with agitation. Blots were again washed 3 times in PBS-T and chemiluminescence generated with Pierce ECL substrate was imaged using an iBright™ instrument (ThermoFisher). Fluorescence was detected using an Odyssey XF imager (LI-COR, Cambridge UK).

### 2.10 Glycosaminoglycan solid-phase binding assays

Heparin from porcine intestinal mucosa (Merck), heparan sulfate (HS) from bovine kidney (Merck), chondroitin sulfate-A (CS) (Merck), were biotinylated as previously described in [46], using the EZ-Link Hydrazine-LC-Biotin kit (Thermofisher). Solid phase binding assays were performed essentially as described previously [47]. Recombinant proteins, either CXCL4, CXCL17 (24-119), SUMO3-CXCL17 (24-119), SUMO3-CXCL17 truncation mutants and SUMO3-tag, were immobilized on 96-well EIA/RIA high binding plates (Corning) in coating buffer 20 mM Na_2_CO_3_ pH 9.6, overnight at RT. Wells were rinsed with 10mM NaOAc, 150 mM NaCl, 2% Tween-20, pH 6.0 and blocked with PBS, 5% BSA at 37°C for 90 min. biotinylated-heparin, HS or CS was added at 1 µg/ml or as otherwise indicated, and bound for 4 hours at RT. Plates were washed with PBS, and the bound GAG was probed by 1:400 streptavidin-HRP (R&D systems), and subsequent incubation with TMB substrate (ThermoFisher) for 10 minutes. Reaction was stopped with 0.2 M H_2_SO_4_, and OD_450nm_ was measured with a SpectraMax i3x instrument (Molecular Devices). Background signals were corrected against blank wells, and data were analysed and fit to non-linear hyperbola where X is concentration, to permit calculation of B_max_ and K_D_ values for GAG interaction with chemokine using Prism 9.2 (GraphPad, San Diego, CA).

### 2.11 Bio-Layer Interferometry (BLI) to assess chemokine: GAG interactions

Real-time assessment of CXCL4 and CXCL17 binding to heparin dp8 and heparan sulfate was assessed via bio-layer interferometry (BLI) on an Octet Red96 system (Sartorius, Goettingen, Germany) using a methodology adapted from a previous study [48]. Streptavidin coated SAX biosensors (Sartorius) were hydrated for 10 mins in assay buffer 10 mM HEPES pH 7.4, 150 mM NaCl, 3 mM EDTA, 0.05% Tween-20, and coated with 0.078 µg/ml biotinylated-heparin dp8 or 7.5 µg/ml biotinylated-heparan sulfate, until 0.5nm and 0.15nm wavelength shifts were detected respectively. SAX sensors were washed in regeneration buffer 0.1 M glycine, 2 M NaCl, 0.1% Tween-20, pH 9.5 and equilibrated in assay buffer. The Octet Red96 system performed a sensor check (20s), baseline reading (60s), association phase (1080s), dissociation phase (1500s), regeneration (30s 3x times) and baseline reading (60s); where reference and GAG-coated SAX sensors sampled 200 µl preparations of chemokine diluted in assay buffer. The binding signal data were recorded at 5 Hz and assessment of the dissociation off-rates of CXCL4 and CXCL17 binding to GAGs were performed in Octet HT 10.0 analysis program. Curves were fitted to a dissociation phase with a Fast 1:1 local model, and maximal responses (nm) and dissociation rate kDis (1/s) were calculated.

### 2.12 Statistical Analyses

All statistical analyses were performed in Prism 9.2, using two-way ANOVA (unless indicated otherwise) with multiple comparisons and Dunnett’s post-test. Statistically significant differences are displayed as p<0.05*, p<0.01**, p<0.001*** and <0.0001****.

## 3 Results

### 3.1 *In silico* characterisation of CXCL17 questions its classification as a chemokine based upon structural features

The CXCL17 gene in primates and rodents encodes a protein of 119 amino acids [7; 17; 49] with considerable homology between the human form ranging from 62.2% identity with the *Mus musculus* CXCL17 orthologue to 98.3% identity with the *Pan troglodytes* CXCL17 orthologue [9]. Post translation, an N-terminal signal peptide is cleaved from the full-length protein CXCL17 (1-119) to liberate mature CXCL17, although the first two publications to describe CXCL17 differ by one amino acid in their predicted cleavage sites [7; 49] despite using the same online prediction software. We used the latest version of SignalP6.0 to predict the likely mature form of CXCL17. The CXCL17 (23-119) species is predicted to be the mature form of human CXCL17 (97.76% likelihood) although the Leu24—Leu119 form of CXCL17 is the form commercially available. This form was used throughout this study unless indicated otherwise and is referred to as CXCL17 (24-119) (Figure 1A). Unique to CXCL17 amongst chemokines is the organisation of the first four cysteines into two CXC motifs, with the first present within an extended N-terminus (Figure 1B). In agreement with a previous report [8], multiple sequence alignments revealed low levels of primary sequence conservation to other CXC chemokines, with only 8.8-18.3% identity. This contrasts with the relatively high levels of structural homology observed amongst the other CXC chemokines, where sequence identity is typically >30% [18], but ranges from 8.87% to 87.67% (data not shown). Subsequent phylogenetic analysis (Figure 1C) suggested CXCL17 is at best, a distant relative of the CXC chemokines with closest homology to CXCL16, another atypical CXC chemokine expressed as a type I membrane protein [10; 50]. The structure of CXCL17 remains uncharacterised experimentally, although folding prediction with the DSC server predicts poor secondary structural homology to CXCL8 (Figure 1D). In contrast, the predicted secondary structure of CXCL8 (Figure 1D) is consistent with experimentally verified structures of the same chemokine (Figure 1E), containing the classical chemokine fold [51]. The 4 cysteine residues of CXCL8 are correctly predicted to form disulfide linkages pairing Cys-7 with Cys-34 (C1-C3) and Cys-9 with Cys-50 (C2-C4) to stabilize the tertiary structure. In contrast, the secondary structure of CXCL17 is predicted to contain 4 α-helices with an absence of β-sheets, inconsistent with the previous structural model generated by Pisabarro and colleagues [7], but supported and expanded upon by Denisov [8].

**Figure 1.**
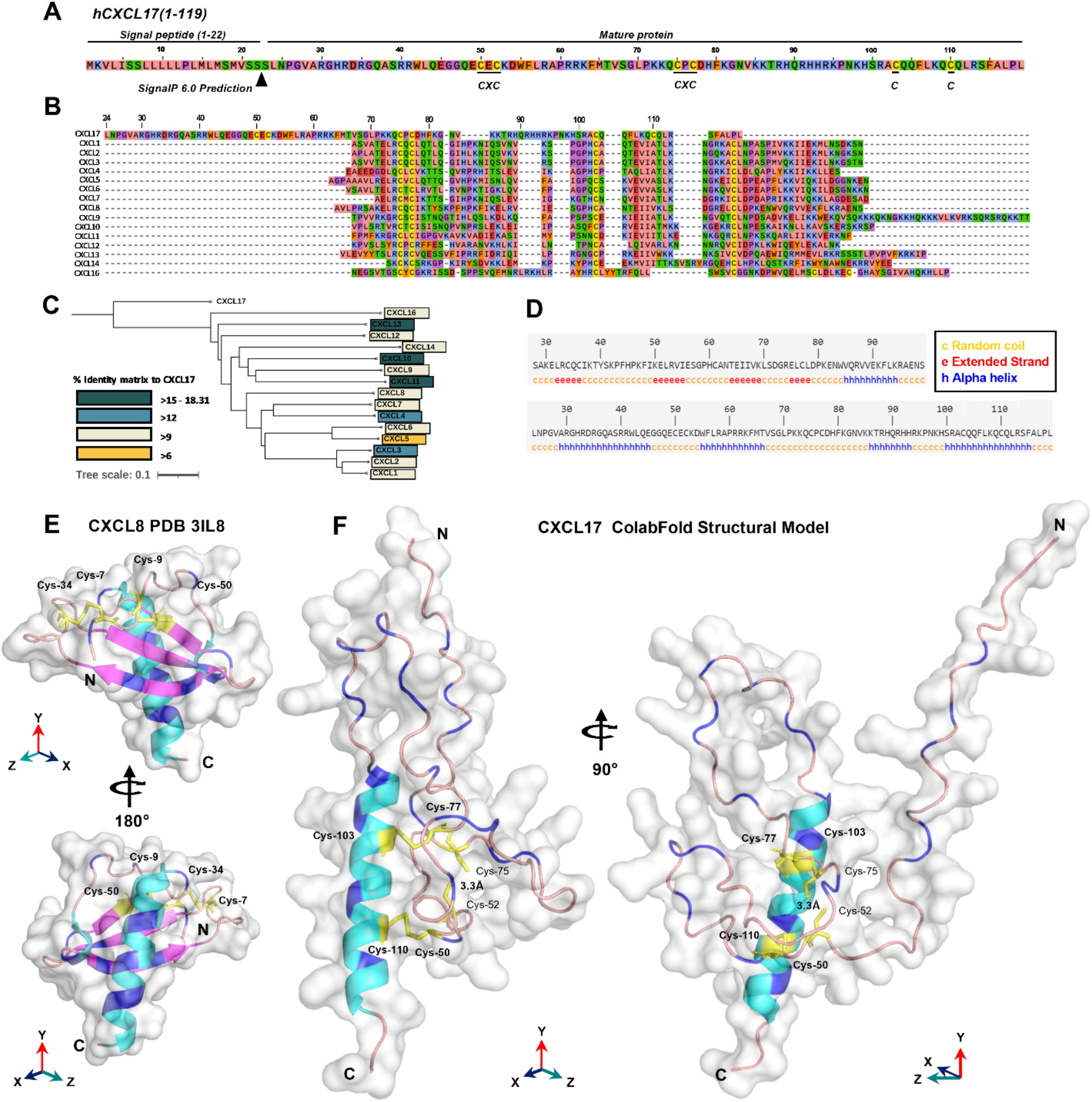
*In silico* analyses raise questions about the classification of CXCL17 as a chemokine. **(A)** The full length amino acid sequence for human CXCL17 (1-119). The predicted signal peptide cleavage site and locations of CXC motifs and cystine-residues are indicated. **(B)** MUSCLE multiple sequence alignment of all human CXC chemokine sequences, aligned against CXCL17 (24-119) using SnapGene. **(C)** Phylogenetic analysis of relationships between all human CXC chemokines. **(D)** DSC protein folding predictions for CXCL8 and CXCL17 (24-119). Regions are marked as random coil (c), extended strand (e) or alpha-helix (a) **(E)** The solved structure of CXCL8. **(F)** Predicted structure of human CXCL17 (24-119) modelled by ColabFold with 48-recycles. E-F were generated in pyMOL with secondary structure indicated (α-helix; cyan, β-sheet; magenta, random coil; salmon pink). The model is displayed with 90° or 180° Y-axis rotation, all cysteines are coloured yellow and annotated, with those in disulphide bonds in bold. Arginine and lysine residues are coloured blue.

In the absence of structural data regarding CXCL17, we performed *in silico* modelling using ColabFold [35]. The results of the highest ranking model generated with AlphaFold2 are shown in Figure 1F (and expanded upon in Figure S1). Structural predictions of CXCL17 folding by AlphaFold2 ColabFold revealed little structural homology to CXCL8 or any other CXC chemokine. The C-terminal α-helix of CXCL17 is predicted to be exposed and is modelled with greater confidence than the rest of the molecule with plDDT residue scores above 70, and low PAE scores (Fig.S1 H-I). Unlike the DSC secondary structure prediction, CXCL17 is not predicted to contain multiple α-helical regions, instead, the majority of the molecule is bundled into loose coiled formations with plDDT scores ranging from 40-65. A 90° rotation on the Y-axis, reveals the N-terminus to be projected away from the core of the molecule. Disulfide bonding is predicted to take place in CXCL17, but contrasts with the C1-C3 and C2-C4 linkages observed in CXCL8 and other CXC chemokines. Disulphide bonds are predicted between Cys-50-Cys-110 (C1-C6) and Cys-77-Cys-103 (C4-C5). No linkage is predicted between Cys-52-Cys-75 (C2-C3), although their close proximity (modelled at 3.3Å) suggests that a third pair of disulfide bonds within CXCL17 is a possibility. Additional structural modelling by C-I-TASSER partly corroborated the structural prediction made by AlphaFold2 ColabFold (Figure 1F), similarly predicting a C-terminal α-helical region, with the remainder of the protein comprised of 6 short α-helices bundled into a compact conformation (Figure S2). Modelling by RoseTTAFold using ColabFold MSA techniques, predicted the presence of 4 α-helices but in an extended conformation, with the C-terminal helix corroborated by the other models also predicted with higher plDDT scores than the rest of the molecule (Figure S3A-C). Due to the limitation of CXCL17 exhibiting low numbers of homologs in the Uniref90 and environmental databases searched on ColabFold, the predictions are of lower confidence than they would otherwise be for other better characterised sequences. To account for this, we also modelled CXCL17 (24-119) using *de novo* folding techniques within AlphaFold (Figure S4) and RoseTTAFold (Figure S3D-F) via ColabFold and again were unable to generate a predicated chemokine fold with both structures comprising of 3-4-α helices interspersed with random coiled regions. In summary, using a complimentary series of modelling techniques, we were able to correctly model CXCL8 from the primary sequence, but were unable to determine a chemokine fold for CXCL17 (24-119).

### 3.2 CXCL17 can form dimers and is not glycosylated post translation

A previous electrophoretic analysis of rat and human CXCL17 expressed by endogenous cells and transfectant cell lines suggested that full length CXCL17 (1-119) undergoes proteolytic cleavage since two bands were observed by western blotting; a larger pro-protein of a little over 20 kDa and a smaller protein running between 6 and 16 kDa [17]. These forms were reported to represent CXCL17 (24-119) and CXCL17 (64-119) although the predicted molecular weights of these molecules (11.3 kDa and 6.6 kDa) do not tally precisely with those observed. We reassessed these findings by expressing the full length ORF of human CXCL17 with a C-terminal His tag in CHO-L-761H cells. Cell lysates were generated and following blotting, were probed with an anti-CXCL17 antiserum. CXCL17 was detected in transfected cells but not in mock-transfected cells running under reducing conditions with estimated molecular weights of around 15 and 30kDa (Figure 2A). Under the same conditions, commercially available human CXCL17 (24-119) which was produced in *E.coli* ran with slightly lower molecular weights of 14 kDa and 28 kDa, resembling the two bands previously reported by Lee and co-workers [17]. Since the recombinant CXCL17 (24-119) had not been exposed to eukaryotic signal peptide cleavage proteases, we hypothesised that these bands represented monomers and dimers of the CXCL17 (24-119) species. Since a property of chemokines is to form dimers and higher-order oligomers as a function of increased concentration [48], we assessed the apparent molecular weights of recombinant CXCL17 (24-119) in the 1.25μM - 10μM range via western blot (Figure 2B). At a concentration of 5μM and above, CXCL17 was observed to form the 28kDa species in an apparent 3:1 ratio of monomer:dimer respectively.

**Figure 2.**
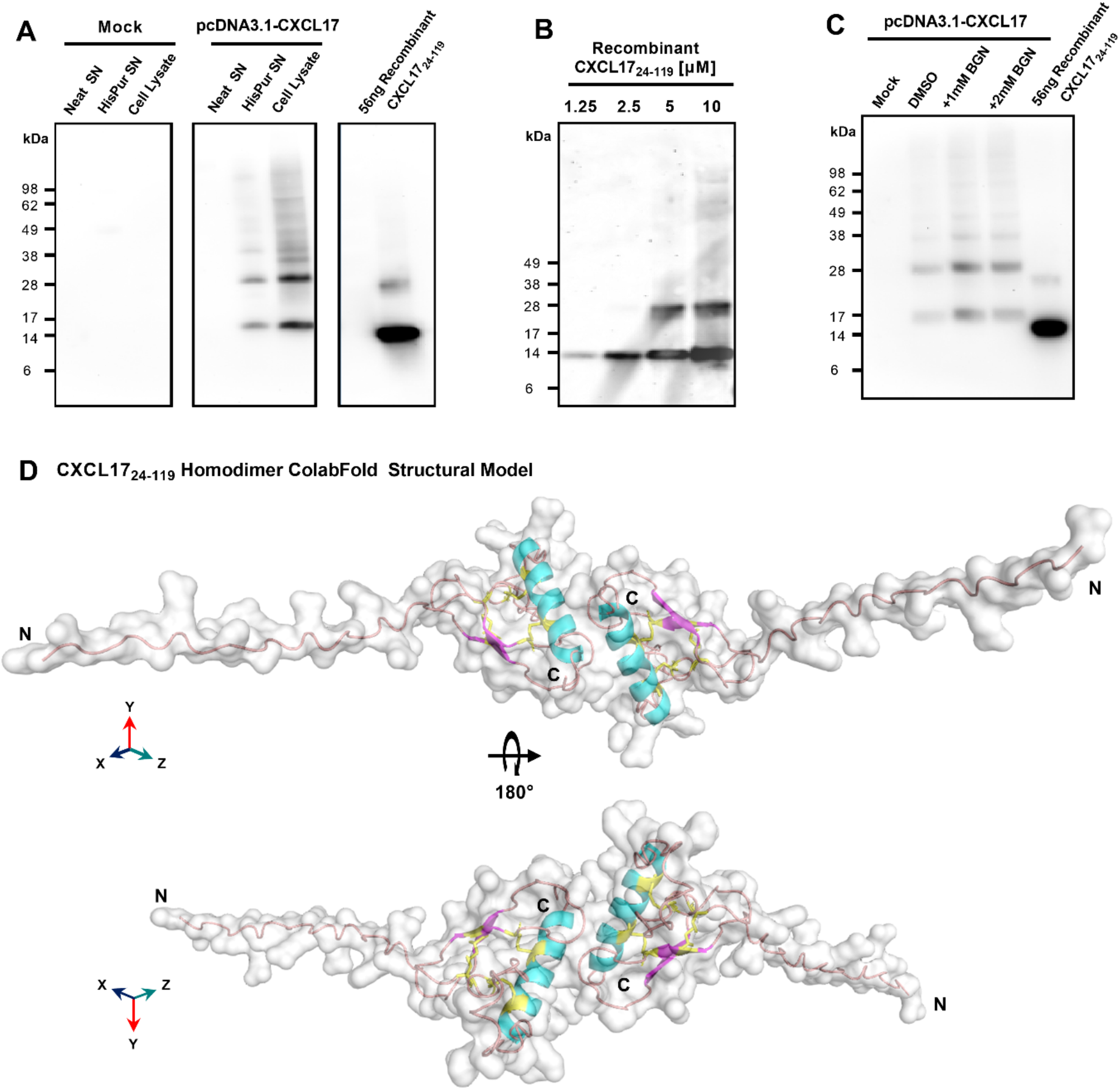
CXCL17 assembles into multimers and is not glycosylated. **(A**) Western blotting of concentrated CHO-761H supernatants following either mock transfection or transfection with pCDNA3.1 containing a C-terminally His-tagged CXCL17 ORF. Recombinant CXCL17 (24-119) was analysed alongside as a control. Data are representative of 3 experiments. (**B**) Western blotting of serial dilutions of bacterially expressed recombinant CXCL17 (24-119). (**C**) *O-* linked glycosylation of CXCL17 in CHO-761H cells was inhibited by culture with the indicated concentrations of Benzyl-GalNac (BGN) or vehicle control (DMSO). All western blots are representative of 3 independent experiments. (**D**) ColabFold structural model of a CXCL17 (24-119) homodimer. Models were generated in pyMOL with molecular surface and secondary structures indicated. The model is rotated 180° in the X-axis, with cysteines coloured yellow and represented as sticks. N and C-termini are annotated with N and C respectively.

Since glycosylation can also influence the apparent molecular weight of proteins, the NetOGlyc-4.0 and NetNGlyc-1.0 servers were used to predict putative glycosylation sites. Seven potential *O-* glycosylation sites but no *N-*linked sites were predicted within the CXCL17 primary sequence. CHO-L-761H cells were transfected with the CXCL17 (1-119) his-tagged construct and *<u>O-</u>*linked glycosylation was inhibited by supplementing cultures with benzyl-2-acetamido-2-deoxy-a-D-galactapyranoside (BGN). Inhibition of *O-*linked glycosylation by BGN revealed no change in apparent molecular mass suggesting that CXCL17 does not undergo post translational glycosylation (Figure 2C).

Since CXCL17 appears to form dimers, a model of CXCL17 (24-119) dimer formation was generated in AlphaFold2 ColabFold (Figure 2D; expanded details in Figure S5). The model predicted that CXCL17 dimerizes along an interface containing their respective exposed C-terminal α-helices, oriented end-to-end. The dimer model has a different pattern of disulphide bond formation than the monomer, with 3-disulfide bonds predicted to form between Cys77-103 (C4-C5), Cys52-110 (C2-6), Cys50-75 (C1-C3). Interestingly, the model also predicts that structural changes take place within each of the constitutive monomers, with the formation of a short region of 2 anti-parallel β-strands, and an extended N-terminus which is projected into space away from the dimer complex.

### 3.3 CXCL17 (24-119) fails to recruit murine splenocytes

CXCL17 has previously been reported to be chemotactic for dendritic cells and monocytes [7] although we and others have struggled to show robust chemotactic activity for monocytes and THP-1 cells using commercially available CXCL17 (24-119) [22; 23]. In modified Boyden chamber assays, recombinant mouse CXCL17 (23-119) was previously reported to attract a subpopulation of splenocytes (CD11b^+^Gr-1^high^ F4/80^-^ cells) isolated from SCID mice in a dose-dependent, pertussis toxin-sensitive manner [19]. We assessed these studies using splenocytes isolated from C57BL/6 mice and a broad concentration range of mouse CXCL17 (23-119). As a positive control chemoattractant 10% FBS was used. In our hands, mouse splenocytes were observed to undertake low levels of chemotaxis in response to FBS, which was statistically significant when compared with basal migration reported as either the percentage of migrating cells or as a chemotactic index (Figure 3A-B). In contrast, exposure of the same splenocytes to a broad concentration range of mouse CXCL17 (23-119) failed to elicit significant levels of chemotaxis and at the highest concentration of 1µM CXCL17 (23-119) resulted in migration at levels significantly lower than those observed in the absence of a stimulus. We therefore conclude that mouse CXCL17 (23-119) is not a major chemoattractant for mouse splenocytes.

**Figure 3.**
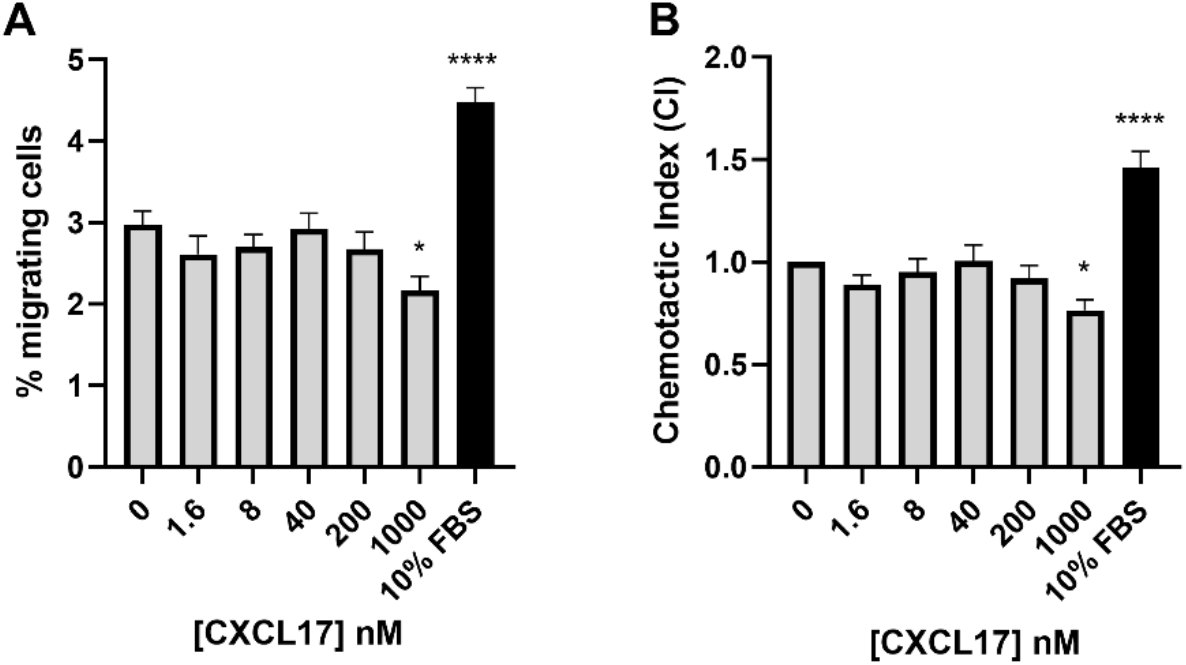
CXCL17 does not induce chemotaxis of murine splenocytes. (**A**) Total % migrating cells are displayed, and (**B**) Chemotactic index (CI) normalized as fold-increase in migration over baseline, of total murine splenocytes in response to serial dilutions of mCXCL17 or to 10% FBS as a positive control (n=4). Data are shown as mean + SEM. One-way ANOVA with multiple comparisons and Dunnett’s post-test was performed and statistical significance is displayed as p>0.05* and p>0.0001****.

### 3.4 CXCL17 is a weak chemoattractant for human neutrophils

To date, no reports of activity for human neutrophils have been described. We therefore investigated the potential for human CXCL17 to recruit freshly isolated human neutrophils, using a real time chemotaxis assay (TAXIScan) [52]. In this system, a chemoattractant gradient is formed by the addition of 1μL of varying concentrations of chemoattractant and the migration of individual cells is tracked microscopically as a function of time. A CXCL8 gradient was used as a positive control.

Human neutrophils exposed to a gradient formed by 1μL of 10nM CXCL8 responded with robust chemotaxis when compared to neutrophils analysed in the absence of a chemoattractant (Figure 4A-B). In contrast, responses to a range of CXCL17 (24-119) gradients revealed little in the way of chemotactic activity until 1μL of 5μM CXCL17 was employed (Figure 4C-F). Analysis of the individual tracks of neutrophils allowed the determination of the velocity, directionality and yFMI parameters (Figure 4G-I). In agreement with the cell tracks, only the parameters of chemotactic responses to 10nM CXCL8 and 5μM CXCL17 were found to be significantly different from those determined in the absence of a chemoattractant. Thus, we conclude that CXCL17 (24-119) has modest chemotactic activity for human neutrophils.

**Figure 4.**
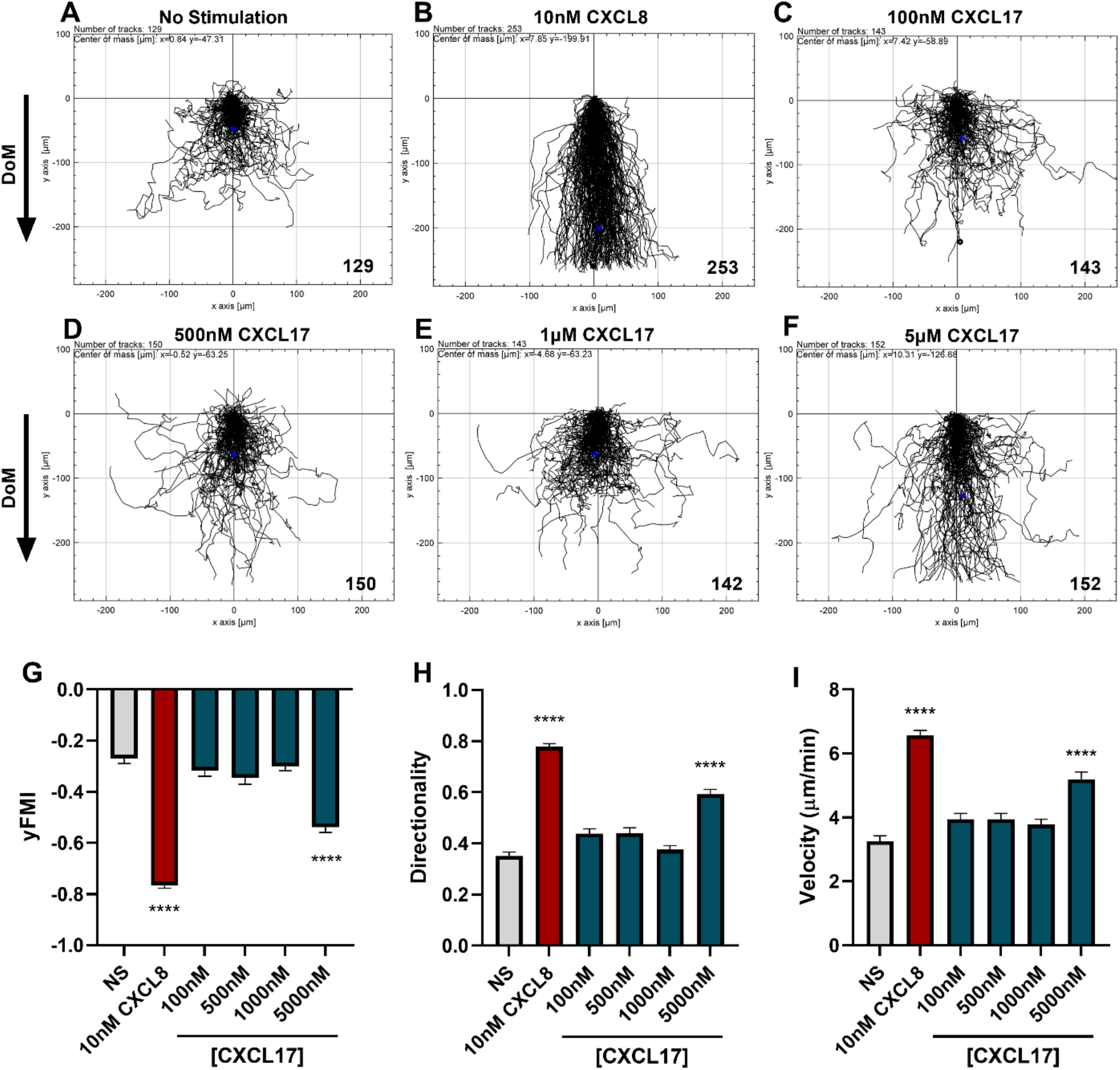
CXCL17 is a comparatively weak chemoattractant for neutrophils. Migration of individual neutrophils were assessed using the EZ TAXIScan real-time chemotaxis assay, in the absence of stimulus (**A**), and after addition of various concentrations of CXCL8 (**B**), or CXCL17 (24-119) (**C-F**). Direction of movement (DoM) is indicated by an arrow. Individual paths were collated, with the total number of cells tracked at each condition indicated in the bottom right of the respective XY-plots. Aggregated chemotaxis from 8 independent donors was analysed and pooled to calculate Y-axis forward migration index (yFMI) (**G**), directionality (**H**) and velocity (**I**) of responses. Data are displayed as mean +SEM, and all statistical analyses were made as comparisons against the no stimulus control. One-Way ANOVA with a Dunnett’s Multiple Comparison test was performed on biological replicates (n=8).

### 3.5 CXCL17 binds glycosaminoglycans with greater capacity than CXCL4

A key requirement for the chemotactic function of many chemokines *in vivo* is the ability to bind to GAGs on the surface of cells [53]. Given the relatively high isoelectric point of CXCL17 we tested the hypothesis that CXCL17 would bind to GAGs using a solid phase binding assay [54]. CXCL4, originally identified due to its heparin binding properties [55], and subsequently characterised as having a low nM affinity for binding heparin, heparan sulfate (HS) and chondroitin sulfate (CS) [48] was used as a positive control (Figure 5A-C). A broad concentration range of immobilized CXCL17 was seen to bind heparin, HS and CS with a significantly greater capacity than equimolar concentrations of CXCL4, as determined by the percentage of maximal binding which in all cases was to either 500 nM or 1 µM CXCL17 coatings. The maximal recovered binding signal of all three GAGs tested was consistently higher when bound to CXCL17 than for CXCL4, indicating that immobilized CXCL17 exhibited a greater capacity for binding GAGs than CXCL4 in this assay. However, care must be taken when interpreting this observation as immobilisation of the chemokine may disrupt typical oligomerisation dynamics that occur when binding to GAGs, and the proteins may have varying adsorption rates to polystyrene plastics which could impact the absolute amount of immobilized chemokine present in this assay.

**Figure 5.**
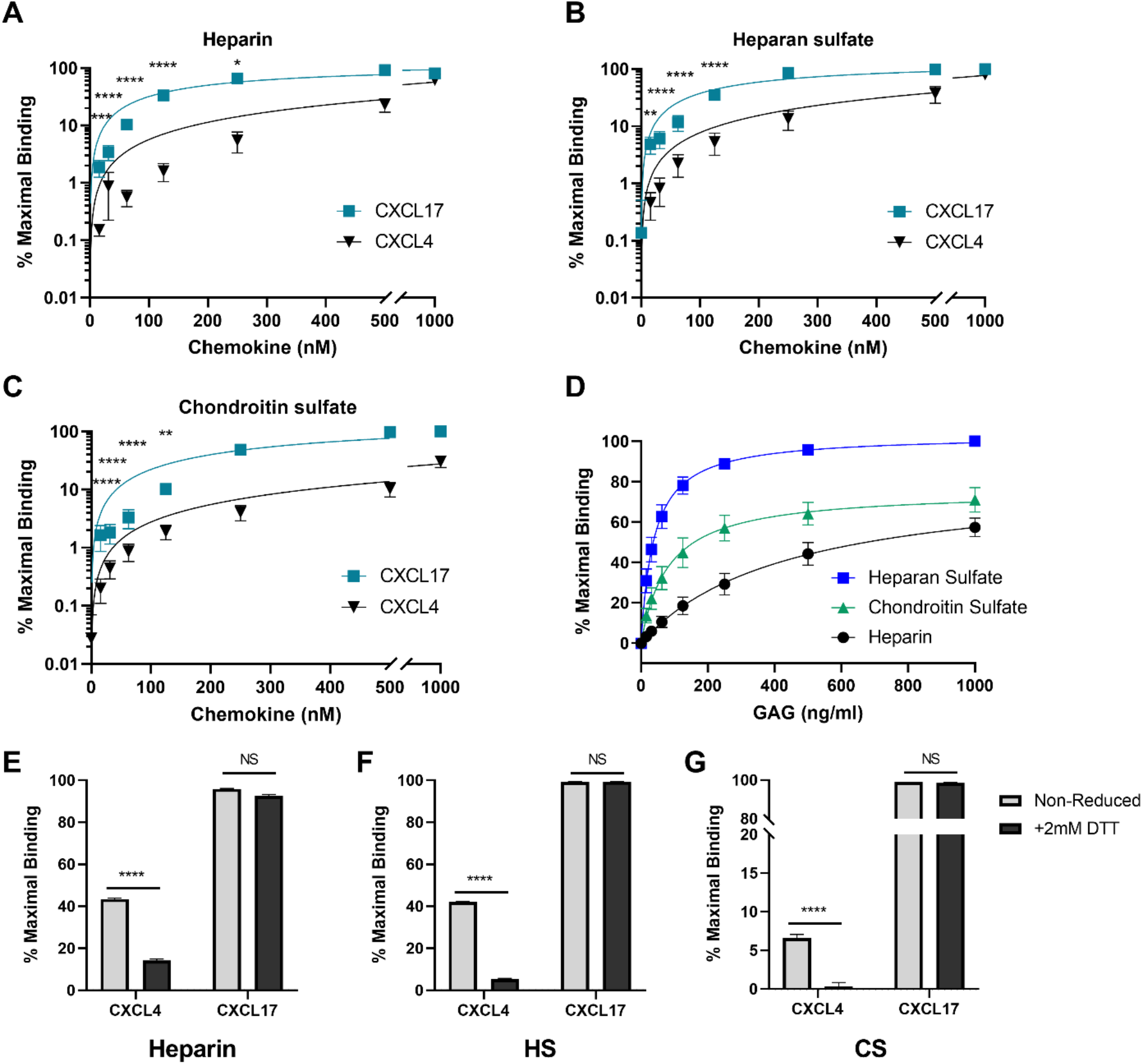
CXCL17 is an efficacious binder of glycosaminoglycans. (**A-C)** The binding of heparin (**A**), HS (**B**) or CS (**C**) by immobilised recombinant CXCL17 (24-119) or CXCL4 (n=4). (**D**) 500nM CXCL17 (24-119) was immobilized, and binding of increasing concentrations of biotinylated heparin, HS and CS was assessed (n=3). (**E-G**) Chemokines were immobilised in both their native state and following reduction with 2mM DTT, after which the binding of 1µg/ml heparin, HS or CS was quantified (n=3). In all cases, data were normalized to the maximum binding signal recorded for the experiment, with data presented as a percentage of this signal and displayed as mean ±SEM.

To directly compare the affinity of binding associations of heparin, HS and CS to CXCL17, a fixed concentration of CXCL17 was immobilised, and varying concentrations of GAG were bound (Figure 5D). Curve fit analysis (Table 1) revealed the calculated *K_D_* values for GAG binding to CXCL17 were in the mid-low nanomolar range. The recovered binding signal was normalised to the maximal signal from the experiment (1000ng/ml HS), which permitted relative comparison between the overall levels of CXCL17 binding capacity for the different GAGs. Immobilized CXCL17 exhibited the greatest capacity for binding HS, and while the binding of the maximum concentration of CS exceeded that of heparin, curve fit analysis revealed the B_max_ value of heparin was higher than CS, indicating that CXCL17 may have a lower affinity, but higher binding capacity for heparin than CS.

**Table 1.**
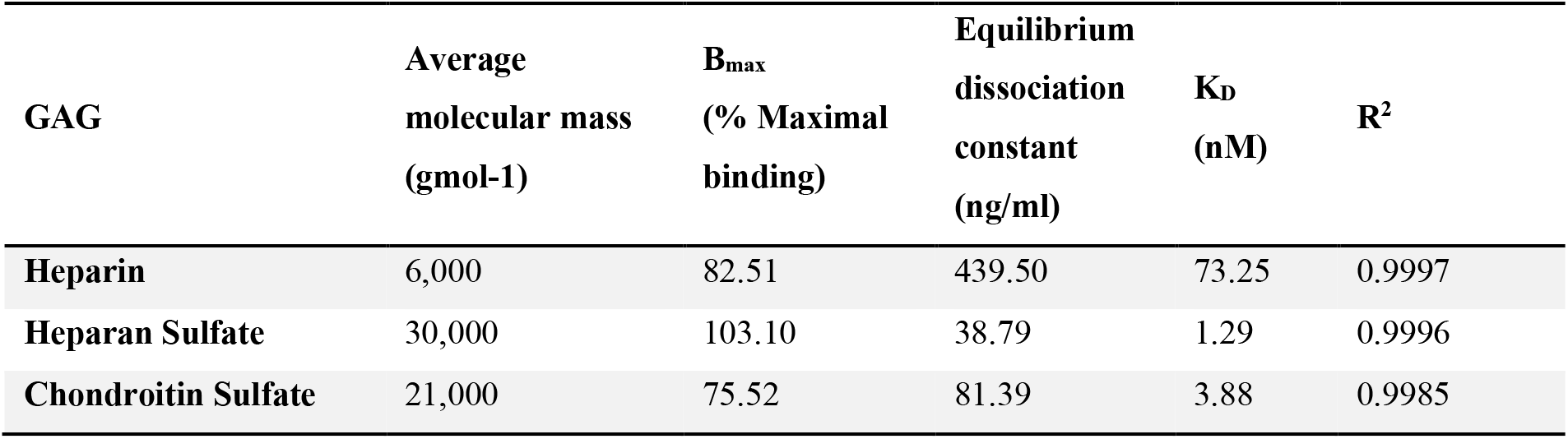
The maximal binding and K_D_ of heparin, heparan sulfate and chondroitin sulfate to immobilized CXCL17 (24-119)_._ Binding associations of heparin, HS and CS to immobilised recombinant CXCL17 (24-119) were calculated using a solid phase binding assay (Fig 4.D). 500nM CXCL17 was immobilized, to which 1000, 500, 250, 125, 62.5, 31.25, 15.625 and 0 ng/ml heparin, HS or CS were bound and quantified at OD_450nm_ by a colorimetric assay (n=3). Curves were fit using GraphPad Prism non-linear fit analysis, and display hyperbola where X is concentration. Best-fit values to the mean data points of 3 independent experiments were calculated for *B_max_* and *K_D_* of each GAG binding to immobilised CXCL17 (24-119). The goodness-of-fit is displayed by the R^2^ values.

| GAG | Average<br>molecular mass<br>(gmol-1) | B<sub>max</sub><br>(% Maximal<br>binding) | Equilibrium<br>dissociation<br>constant<br>(ng/ml) | K<sub>D</sub><br>(nM) | R<sup>2</sup> |
| --- | --- | --- | --- | --- | --- |
| Heparin | 6,000 | 82.51 | 439.50 | 73.25 | 0.9997 |
| Heparan Sulfate | 30,000 | 103.10 | 38.79 | 1.29 | 0.9996 |
| Chondroitin Sulfate | 21,000 | 75.52 | 81.39 | 3.88 | 0.9985 |

We also assessed the extent to which the CXCL4 and CXCL17 tertiary structure contributes to the GAG-binding activity by reducing intermolecular disulphide bonds within CXCL17 and CXCL4. These proteins were then immobilised and chemokine binding to heparin, HS and CS assessed (Figure 5E-G). Reduction significantly reduced the binding of all three GAGs to CXCL4, but had negligible impact on GAG binding to CXCL17, suggesting that the interactions of CXCL4 and CXCL17 with GAGs have different structural requirements.

### 3.6 Bio-layer interferometry reveals CXCL17 and CXCL4 have comparable dissociation-rates when binding heparin and heparan sulfate

The GAG binding interactions of CXCL17 and CXCL4 were further investigated by bio-layer interferometry (BLI) [56; 57]. In this assay, the dynamic binding interactions between sensors coated with biotinylated heparin dp8, or HS and soluble chemokine analyte were quantified by optical wavelength shift during association and dissociation stages of binding. Heparin had a greater capacity to bind CXCL17 than CXCL4 at all concentrations of chemokine tested (Figure 6A). Similarly, the binding capacity of HS for CXCL4 increased proportionally with increasing concentrations to higher total levels than CXCL17 at equivalent concentrations (Figure 6B), indicating that the dynamics of chemokine multimerization differ in a GAG-dependent manner. The binding of 250 nM CXCL17 to heparin and HS was characterised by a biphasic association curve for which a global, 1:1 model “Global full” could not be confidently applied, where for the first 100 seconds of interaction, the rate of association was rapid then decelerated until ∼250 seconds where it reached its maximal association. Such deviations from pseudo-first order binding are typically observed as a feature of mass transport limitation (MTL), or may instead be explained due to surface ligand heterogeneity and the presence of multivalent attachment sites for the analyte due to oligomer formation [58; 59]. MTL effects occur more frequently when binding to dense heterogenous ligands like GAGs, which influence local analyte diffusion to and from the bulk, but here we know that both CXCL17 and CXCL4, exhibiting multimer forming capabilities, may simultaneously interact with the surface via multiple interaction sites. This biphasic association was only apparent when 250nM CXCL17 was used and may suggest differing affinities of interaction for CXCL17 with GAGs, and between CXCL17 monomers during multimer assembly. When CXCL17 was diluted to 62.5nM or below, the biphasic curve of binding to heparin and HS was lost, perhaps since the rate of CXCL17 dimer formation was diminished at lower concentrations of chemokine. For CXCL4, no biphasic curves were observed, which suggests that CXCL4 binds heparin and HS with approximately the same affinity with which it forms multimers, with both events occurring at physiologically relevant concentrations of chemokine.

**Figure 6.**
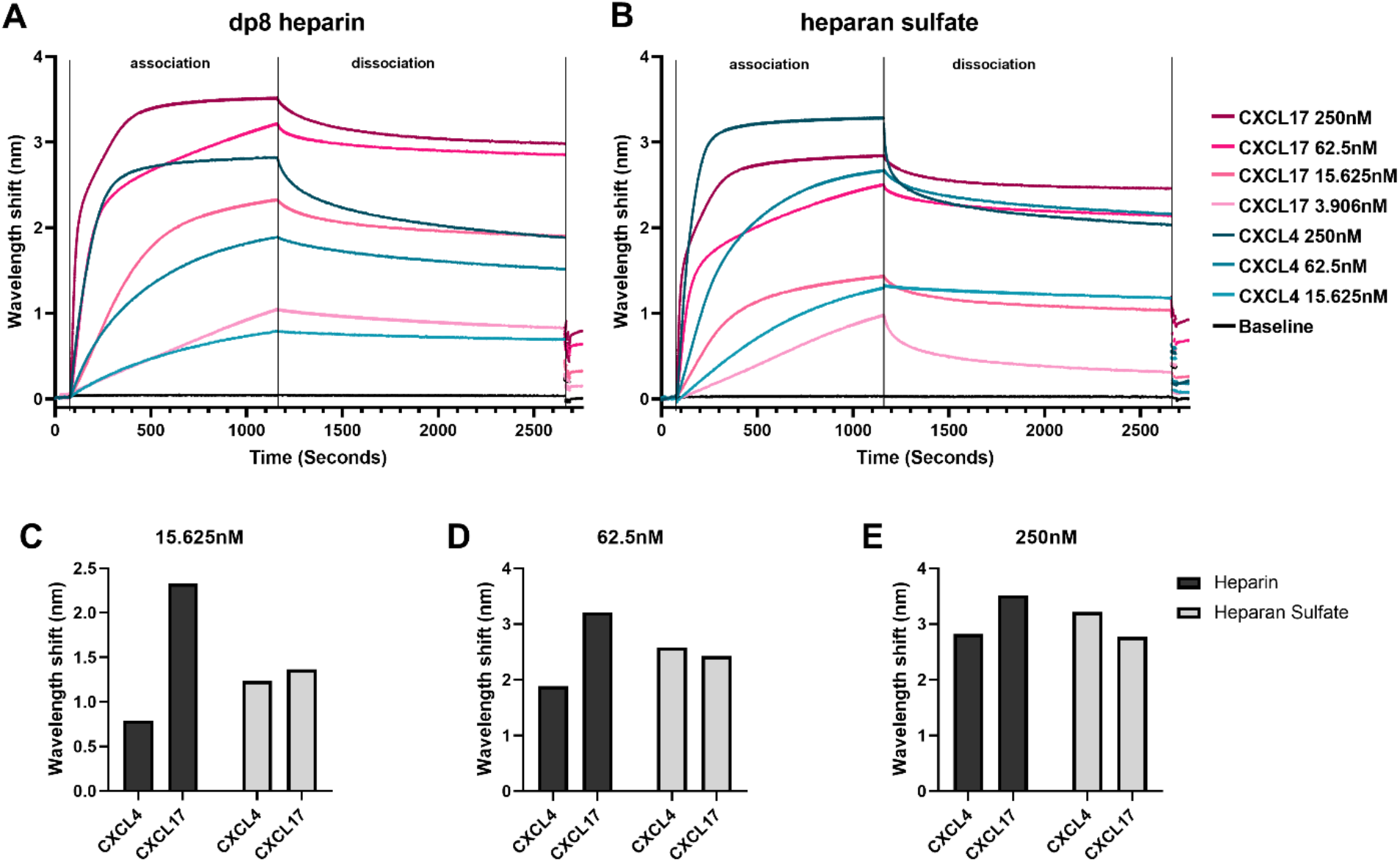
CXCL4 and CXCL17 bind heparin and heparan sulfate with high capacities. Real-time assessment of CXCL4 and CXCL17 (24-119) binding to immobilized heparin (dp8) and HS as assessed by bio-layer interferometry (BLI). Sensorgrams illustrate the binding associations between streptavidin immobilized biotinylated-heparin (**A**), or HS (**B**), and various equimolar concentrations of CXCL4 and CXCL17. Prepared solutions of chemokine were sampled with BLI (Octet) streptavidin coated sensors containing bound GAG, for a 1080 second association phase, and a 1200 second dissociation phase. The experiment was performed in singlicate with data displayed as wavelength shift (nm). (**C-E**) The extent of molecular association of CXCL4 or CXCL17 with heparin or HS at increasing concentrations of chemokine. Measurements were taken of induced wavelength shift (nm), with maximal recorded wavelength shifts for 15.625nM (**C**), 62.5nM (**D**), and 250nM (**E**) of chemokine.

Due to the inability to fit a global, 1:1 model, we opted to compare the dissociation curves for CXCL4 and CXCL17 for which high quality local curve fits could be applied, and for which previous studies have demonstrated a relationship between slow rates of dissociation and high binding affinity for CXCL4 [60]. The dissociation rate for CXCL4 vs HS or heparin, and for CXCL17 vs heparin, decreased alongside the analyte concentration (Table 2). Such an observation is characteristic of MTL, but may also occur when a sub-population of analyte oligomers is bound simultaneously to the surface via multiple interaction sites, with this sub-population exhibiting slower dissociation kinetics since both attachments must be broken simultaneously to permit free dissociation [58]. Conversely, the dissociation rate for CXCL17 binding to HS was not dose dependent in the same manner and would therefore not support these curve deviations being MTL mediated. Together, these findings suggest that when chemokine concentration is highest, the rate of dissociation from the GAGs is fastest. We may speculate this is partly driven by chemokine dissociation from clustered multimer complexes on the bound GAG, where the affinity for binding the multimer complex is lower than the affinity for binding GAG. Together these observations suggest concentration dependent CXCL17 multimers may play a role in mediating dynamic GAG interactions.

**Table 2.**
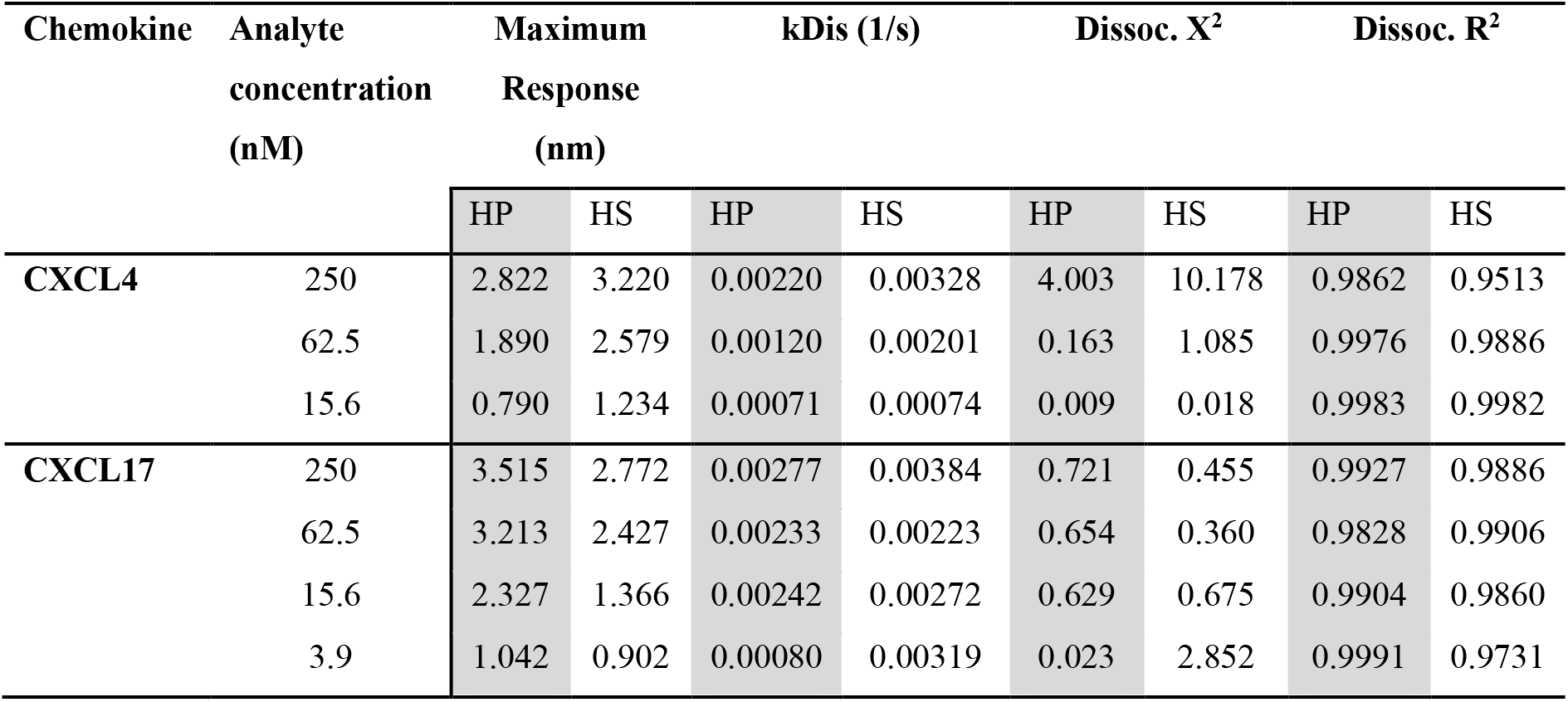
CXCL4 and CXCL17 bind GAGs with high capacity and with comparable dissociation rates. The dissociation rates (kDis) of CXCL4 and CXCL17, from heparin (dp8) and HS were calculated using curve fit analysis software from Octet (Sartorius). Local individual curve fits were performed to the dissociation phase of the BLI sensorgrams using a fast 1:1 local model and were used to calculate the kDis (1/s). The quality of the data fitting is reflected by the *X^2^* value, and the goodness-of-fit is expressed by the *R^2^* value. Data display values calculated from individual curve fits performed in singlicate.

| Chemokine | Analyte<br>concentration<br>(nM) | Maximum<br>Response<br>(nm) |  | kDis (1/s) |  | Dissoc. X<sup>2</sup> |  | Dissoc. R<sup>2</sup> |  |
| --- | --- | --- | --- | --- | --- | --- | --- | --- | --- |
|  |  | HP | HS | HP | HS | HP | HS | HP | HS |
| CXCL4 | 250 | 2.822 | 3.220 | 0.00220 | 0.00328 | 4.003 | 10.178 | 0.9862 | 0.9513 |
|  | 62.5 | 1.890 | 2.579 | 0.00120 | 0.00201 | 0.163 | 1.085 | 0.9976 | 0.9886 |
|  | 15.6 | 0.790 | 1.234 | 0.00071 | 0.00074 | 0.009 | 0.018 | 0.9983 | 0.9982 |
| CXCL17 | 250 | 3.515 | 2.772 | 0.00277 | 0.00384 | 0.721 | 0.455 | 0.9927 | 0.9886 |
|  | 62.5 | 3.213 | 2.427 | 0.00233 | 0.00223 | 0.654 | 0.360 | 0.9828 | 0.9906 |
|  | 15.6 | 2.327 | 1.366 | 0.00242 | 0.00272 | 0.629 | 0.675 | 0.9904 | 0.9860 |
|  | 3.9 | 1.042 | 0.902 | 0.00080 | 0.00319 | 0.023 | 2.852 | 0.9991 | 0.9731 |

Additionally, it was observed that higher concentrations of CXCL17 failed to completely dissociate from the immobilized heparin or HS during the wash step performed between experimental replicates (Figure 6 A-B). While this observation further supports the notion of multivalent binding between CXCL17 and immobilised GAG, this retention of binding signal introduced a confounding artefact for subsequent experimental duplicates. Therefore, only the data acquired from the first unaffected replicate was incorporated into this study.

The total extent of chemokine-GAG binding associations was estimated by the amount of optical wavelength shift (nm) on the BLI sensor surface, which is proportional to the amount of analyte mass bound to the GAG-coated sensor. Taking different concentrations of CXCL4 and CXCL17 (15.625, 62.5 and 250 nM) and comparing the maximal wavelength shift at the association phase end-point, the relative amount of chemokine accumulation on the GAG was inferred (Figure 6C-E). In keeping with our solid-phase binding assays (Figure 5), we saw that on heparin, CXCL17 accumulated at higher levels than CXCL4, suggesting that in this assay with immobilized GAGs, heparin has a higher capacity for binding CXCL17 than CXCL4 (Figure 6C-E). On HS, the total binding capacity for CXCL4 and CXCL17 was approximately equal at 15.625 nM (Figure 6C) but was greater for CXCL4 at concentrations above this (Figure 6D-E). All in all, the BLI assays determined that under dynamic conditions containing free chemokine and immobilized GAG, CXCL17 exhibits selectively greater accumulation on heparin while CXCL4 accumulated to a greater total extent on HS.

### 3.7 Truncation of CXCL17 identifies GAG binding motifs within the C-terminus

GAG binding motifs encoded within the primary structures of several chemokines have been studied by several groups and typically rely on the juxtaposition of amino acids with basic side chains such as lysine and arginine. This is exemplified by the BBXB and BBBXXB motifs where B represents a basic amino acid [61]. Analysis of the CXCL17 (24-119) sequence revealed several putative GAG binding motifs as highlighted in Figure 7A. We therefore carried out a program of mutagenesis to identify residues involved in GAG binding. Four truncation mutants (Δ20, Δ40, Δ60 and Δ80) were generated in which the C-terminus of CXCL17 was progressively truncated by 20 amino acids.

**Figure 7.**
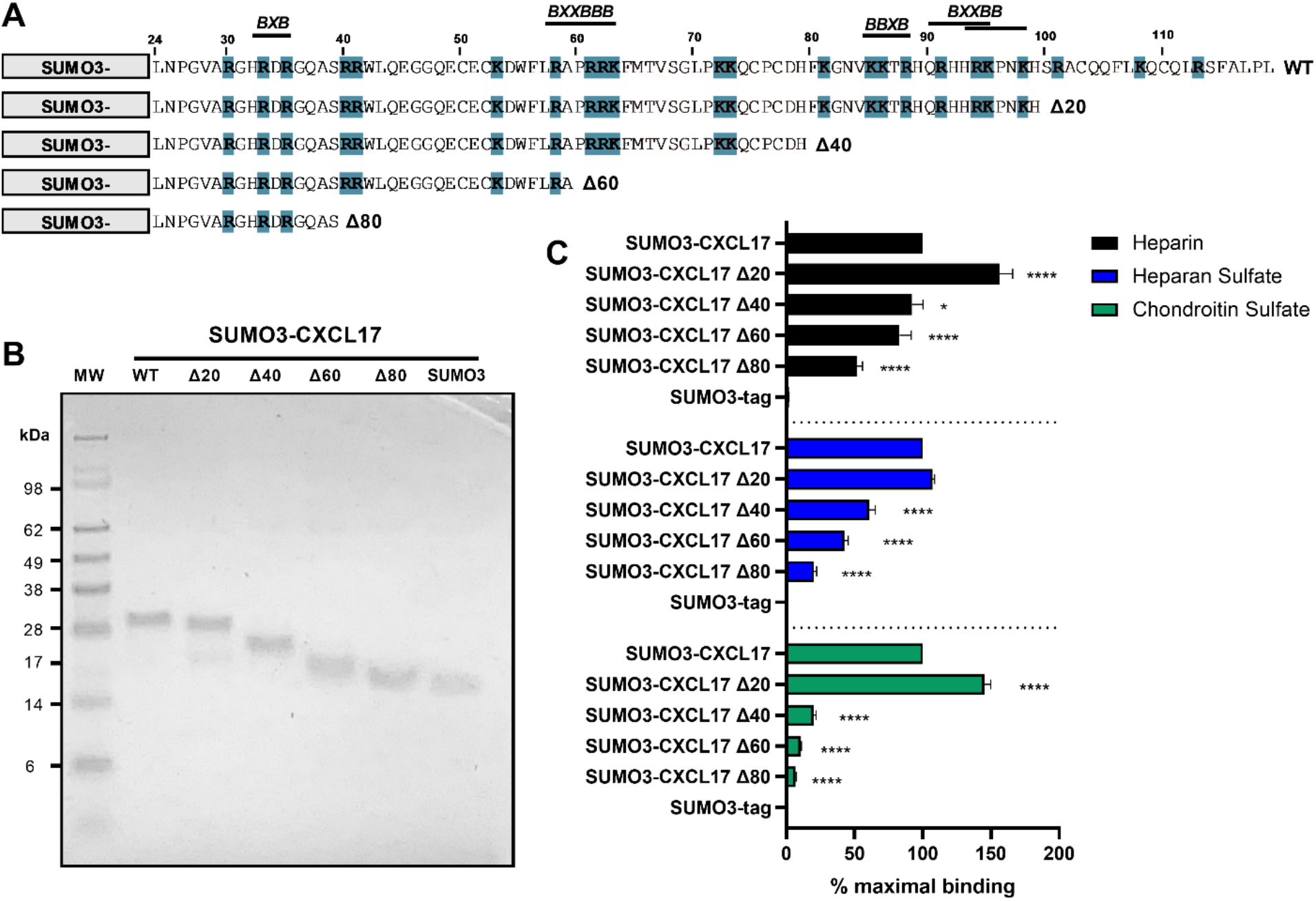
The extreme C-terminus of CXCL17 occludes a cryptic GAG-binding domain. (**A**) Schematic representation of C-terminal truncations of SUMO3-CXCL17 (24-119) generated. Lysine and arginine residues are highlighted blue and putative GAG binding regions indicated, where B denotes a basic residue and X denotes any residue. (**B**) Recombinant SUMO3-CXCL17 truncations as resolved by reducing SDS-PAGE. (**C**) Binding interactions of GAGs to immobilised SUMO3-CXCL17 truncation mutants as assessed using a solid phase binding assay. Chemokines were immobilised and the binding of heparin, HS and CS was quantified, with the % maximal binding normalized to that of the full length SUMO3-CXCL17 (24-119). Data are displayed as mean +SEM (n=4).

Proteins were expressed as N-terminal fusions with the partner protein SUMO3 which we and others have previously used to improve the stability and yield of recombinant chemokine [40; 41].

Typically, in this system the SUMO3 portion is removed post expression via enzymatic cleavage with *UlpI* to yield the mature protein (Figure S6). However, given that some truncation mutants were short and difficult to resolve via SDS-PAGE, we opted to retain the N-terminal SUMO3 fusion partner having first shown that SUMO3 does not bind to GAGs nor does it interfere with the ability of SUMO3-CXCL17 to bind to GAGs (Figure S7). Constructs encoding SUMO3-CXCL17 (24-119) and the CXCL17 truncation mutants were expressed and purified to homogeneity (Figure 7B). Subsequently, they were assessed for their ability to bind a fixed concentration of heparin, HS or CS using a solid-phase GAG-binding assay (Figure 7C). Binding to all three GAGs was improved by removal of the first 20 C-terminal residues (Δ20 construct). Specifically, binding of the Δ20 construct to heparin and CS, significantly increased to more than 140% of that observed with the parent SUMO3-CXCL17 (24-119). Removal of a further 20 residues (Δ40 construct), resulted in significant decreases in binding to all three GAGs when compared with SUMO3-CXCL17 (24-119) with the greatest reduction observed in the binding of CS. Consecutive deletion of a further 20 amino acids (Δ60 and Δ80 constructs) reduced the binding of all three GAGs, with heparin binding the most resilient to truncation, retaining around 45% of the heparin binding capacity of SUMO3-CXCL17 (24-119) even when 80 C-terminal residues had been removed.

### 3.8 CXCL17 inhibits CXCL8-mediated chemotaxis of CXCR1-Ba/F3 transfectants

Interactions with GAGs on the neutrophil cell surface are essential for responses to the chemokine CXCL8 as highlighted by the pathogen *Streptococcus pyogenes.* Invasive strains of this bacterium produce an enzyme known as SpyCEP which cleaves the major GAG binding domain from CXCL8, rendering the chemokine impotent [44]. We therefore postulated that CXCL17 may be able to inhibit chemotactic responses to CXCL8 by disrupting GAG binding. In modified Boyden chamber assays using a Ba/F3 cell line expressing CXCR1, a robust characteristic bell-shaped dose-response to CXCL8 was observed with CXCL17 inactive over the same concentration range (Figure 8A). No chemotaxis of parental Ba/F3 cells in response to a range of CXCL8 or CXCL17 concentrations was detected (data not shown). Employing a sub-optimal concentration of CXCL8 (0.1nM) in the presence or absence of increasing concentrations of CXCL17, we found that concentrations of CXCL17 between 1nM and 1μM significantly inhibited chemotactic responses to CXCL8, with the uppermost CXCL17 concentration reducing migration to basal levels (Figure 8B). This suggests that whilst having modest chemotactic activity for neutrophils, CXCL17 has the potential to modulate the responses of potent neutrophil-recruiting chemokines.

**Figure 8.**
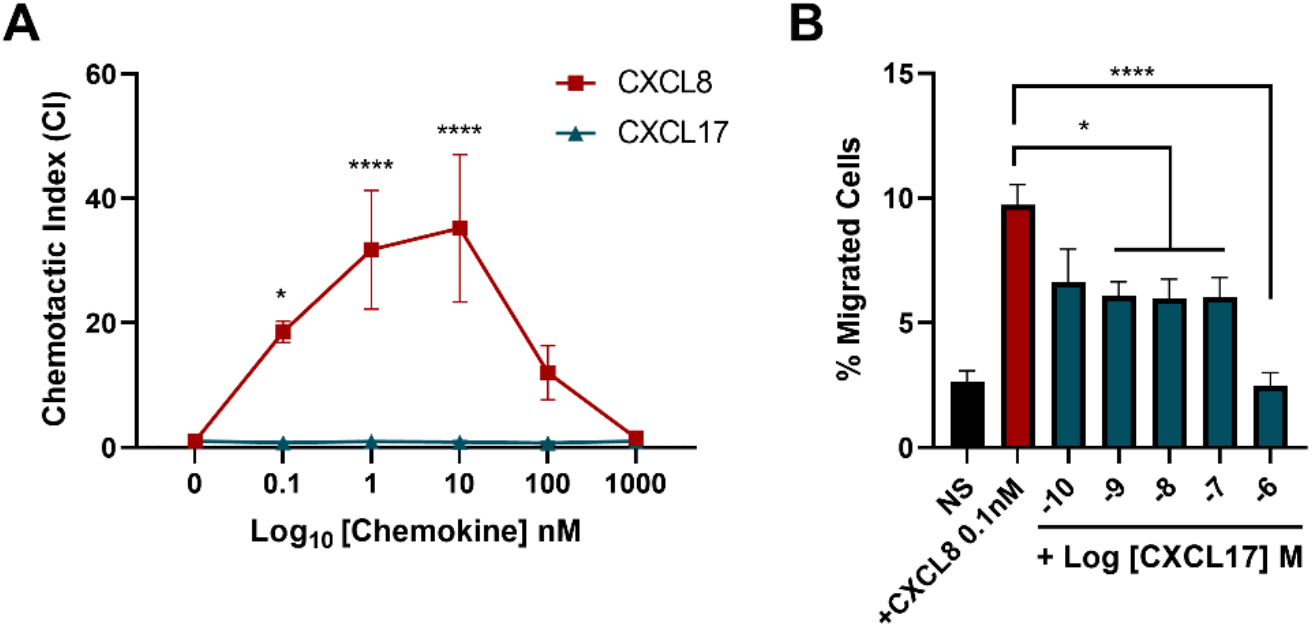
High concentrations of CXCL17 can antagonise the chemotactic responses of CXCR1 transfectants to CXCL8. (**A**) Chemotactic responses of CXCR1 expressing Ba/F3 cells to increasing concentrations of CXCL8 and CXCL17 (24-119). Two-way ANOVA with multiple comparisons and Fishers LSD test was performed and statistical significance is displayed as p>0.05* or p>0.0001****. (**B**) Chemotactic responses of CXCR1-Ba/F3 cells to CXCL8 in the presence of absence of increasing concentrations of CXCL17 (24-119). Data are displayed as the mean +SEM of three independent experiments.

## 4 Discussion

CXCL17 remains a poorly characterised protein with some seemingly contradictory reports published in the literature. In this study, we showed that CXCL17 has some but not all of the properties of a chemokine. CXCL17 can form dimers like many chemokines and has relatively modest chemotactic activity, although it appears unlikely to adopt a chemokine fold. We also showed that CXCL17 is an efficacious binder to GAGs via conserved basic residues within the C-terminus, and that GAG binding may facilitate an antimicrobial activity at mucosal surfaces in addition to modulating the function of other chemokines dependent upon interactions with GAGs for their signalling.

To date, the structure of CXCL17 has not been experimentally verified, with *in silico* modelling generating contradictory reports [7; 8; 9; 62; 63]. Using multiple fold prediction programs and structural modelling algorithms, our methodologies were consistent in predicting that CXCL17 is unlikely to adopt a chemokine fold. This aligns with the inability of BLAST and Pfam analysis of the CXCL17 sequence to find statistically significant structural homologs with any other characterised protein, as confirmed in an earlier report [7]. The structure of CXCL17 (Q6UXB2) has been predicted by AlphaFold (Deepmind, EMBL-EBI) and found to lack structural homology with chemokines [64]. However, the utility of that model may be limited due to the inclusion of the predicted N-terminal signal peptide. Using AlphaFold2 and RoseTTAFold on ColabFold we therefore predicted the structure of the mature CXCL17 (24-119). Acknowledging the limitations of *in silico* structural modelling for proteins with few sequence homologs in reference libraries, we performed MMSeqs2 MSA of CXCL17 on 29.3 million cluster consensus sequences [35], to give the best chance of finding high quality alignments to inform folding. ColabFold identified ∼130 homologous sequences and predicted folding with AlphaFold2 (Figure 1F, S1) or RoseTTAFold (Figure S3A-C). While the structures generated from both methods are not in agreement, they both notably lack a chemokine fold. Due to the relatively low numbers of CXCL17 homologs in reference libraries, we additionally performed *de novo* folding in AlphaFold2 (Figure S4) and RoseTTAFold (Figure S3D-E) whereby the MSA step is bypassed. This approach has been reported to increase folding accuracy for proteins lacking homologs [35]. A higher confidence model was generated in AlphaFold2 using this methodology (Figure S4), as determined by PAE and plDDT scores. Inspection of the resulting *de novo* RoseTTAFold model, produced a CXCL17 structure which again lacked a chemokine-like fold (Figure S3 D-F).

The lack of predicted chemokine-like homology produced by ColabFold was corroborated by C-I-TASSER modelling (Figure S2) and supported previous I-TASSER models reported in Giblin and Pease., [9], Sun *et al.,* [62], and Nijja *et al.,* [63], although in the latter two articles, no discussion of the lack of a chemokine-like structure for CXCL17 was made by the authors. The relatively low confidence of the C-I-TASSER models indicates low-significance of threading template alignments due to a lack of experimentally verified homologs, which itself, speaks to the low conservation between CXCL17 and all other chemokines. Perhaps the generation of several non-chemokine like structural predictions for CXCL17 is unsurprising, given that the N-terminally extended region of CXCL17 with its two CXC motifs is not conducive to alignment with other CXC-chemokines. One likely flaw in the initial prediction of Pisabarro *et al.,* is the usage of a limited quality of homologs (7,950 PDB library entries) of which the 6^th^ and 12^th^ ranked proteins (CCL3: 16.2% identity) and CXCL8 (14.3% identity) were used to generate a new fold library containing all known CXCL8 mutants at that time [7]. Notably, CXCL17 was postulated to form a chemokine fold resembling that of the CXCL8 Glu38Cys/Cys50Ala mutant (1ICW) which was deliberately engineered to modify the disulphide bonding of the chemokine with deleterious effects on receptor binding [65]. Significantly, by their own confidence criteria, other proteins in the library with higher homology were ignored, such as the RXR-α DNA-binding domain (1dsz.B) for which the solved structure comprises of a zinc-stabilized bundle of 3 α-helices with a short length of antiparallel β-sheet [66]. The helical portions of the RXR-α DNA-binding domain protein are in fact more consistent with *in silico* generated models of CXCL17 presented here. In conclusion, we find no evidence to support the likelihood of CXCL17 adopting a chemokine-like fold. Experimental verification of the CXCL17 structure by 2D-NMR or crystallography is required to provide a definitive answer to the structural basis for classifying CXCL17 as a chemokine.

Chemokines are noted for their propensity to form higher-order oligomers as a function of increasing concentration [67]. Early efforts in the field were made by groups to generate mutant forms of chemokines less prone to oligomerisation, for example the BB-10010 form of CCL3. By virtue of a Asp26-Ala mutation, the BB-10010 variant of CCL3 has greatly increased solubility, facilitating its clinical evaluation as a mobilising agent of stem cells [68]. Here, we show that recombinant CXCL17 expressed in either prokaryotic or eukaryotic systems readily forms dimers as revealed by the apparent molecular weight on Western blot. ColabFold modelling suggests that the CXCL17 dimer interface is likely to be at their C-termini, although as with the CXCL17 monomer, these models require structural verification. A wider point illuminated by our observation is that a reinterpretation of a previous study regarding the potential for proteolytic cleavage of CXCL17 may be in order. Lee *et al* previously reported that CXCL17 expressed by cells from the rat stomach ran as proteins of around 8 kDa and 22 kDa as visualised by SDS-PAGE/Western blotting [17]. These data were corroborated using HEK-293T cells expressing human CXCL17 (1-119). The authors of that study interpreted their observation as being evident of the post-translational cleavage of a larger CXCL17 pro-protein into a smaller CXCL17 (64-119) form. They suggested that three basic residues between the second and third cysteines of CXCL17 was the likely site of cleavage and went on to show that expression of a mutant CXCL17 (1-119) in which the tribasic motif was mutated to alanine resulted in expression only of the 22kDa species. A more feasible explanation in our opinion, is that the two species of CXCL17 represents monomer and dimer variants of CXCL17 (24-119). Such an interpretation would also fit better with the apparent molecule weights of the CXCL17 forms on SDS-PAGE. The apparent failure of the Lys61-Arg63 triple alanine mutant of CXCL17 (1-119) to undergo cleavage may be explained by the possibility that the mutation tips the balance in favour of CXCL17 dimer formation [16].

In a similar fashion, although CXCL17 was originally described as a dendritic and monocytic cell chemoattractant protein [7], supporting data from other groups describing chemoattractant properties for CXCL17 is mixed. Notably, GPR35 was previously postulated to be a receptor for CXCL17 [21] which has been questioned by other groups, including our own [22; 23]. A study by Matsui and colleagues suggested that mouse CXCL17 (23-117) was chemotactic for splenocytes with optimum migration observed at a concentration around 20nM, an order of magnitude more potent than the previously described activity of human CXCL17 for monocytes and dendritic cells [7; 19]. However, we were unable to reproduce these findings, with no significant splenocyte chemotaxis to CXCL17 observed above the basal response. We note that Matsui and colleagues used splenocytes isolated from SCID mice whereas our splenocytes were isolated from C57BL/6, so perhaps this biological difference may explain our divergent findings.

Using the TAXIScan system, we observed that human CXCL17 (24-119) was chemotactic for human neutrophils which to our knowledge, is the first report of CXCL17 recruiting this leukocyte population. However, this activity was weak, with significant chemotaxis only observed at CXCL17 concentrations 500-fold higher than the CXCL8 positive control. The CXCL17 (24-119) form has a greatly extended N-terminus when compared to other CXC chemokines which is hard to reconcile with current models of chemokine receptor activation [69; 70]. In these models, a relatively short N-terminus inserts into the helical bundle to induce the conformation changes required for productive G protein coupling. The generation of a CXCL17 (64-119) form postulated by Lee and colleagues, although having four cysteine residues like most other CXC chemokines is unlikely to adopt a chemokine-like fold, therefore any chemotaxis is likely to be mediated via a mechanism atypical of CXC chemokines [70]. Perhaps tellingly, the functional validation of commercially available recombinant CXCL17 (24-119) from a variety of sources does not report chemotactic activity for leukocytes but the ability to induce VEGF expression in mouse endothelial cells [49]. This leads us to conclude that the primary biological function of CXCL17 may not be to promote leukocyte chemotaxis.

Using solid-phase and BLI-based assays, we report that CXCL17 binds with high affinity to a variety of GAGs, as may be predicted given the high proportion of basic residues within the primary sequence. Direct comparisons with the well-characterised GAG-binding chemokine CXCL4 [48] found that CXCL17 bound to GAGs with comparably high capacity, often exceeding that of CXCL4. Additionally, CXCL4 and CXCL17 had similar dissociation rates from heparin and heparan sulfate which may be indicative of comparable binding affinities. In contrast to CXCL4, reduction of the disulphide bonds within CXCL17 had negligible effect upon GAG binding, suggesting that tertiary conformation plays little role in CXCL17-GAG binding. Serial truncation of CXCL17 revealed that a BBXB and BXXBB motif conserved between residues 85-98 appeared to be a major determinant for binding to heparan sulfate and chondroitin sulfate. Taken together with the observation that CXCL17 can form dimers, it suggests that CXCL17 dimers may form on GAGs as a function of increasing CXCL17 concentration. So what might be the function of GAG-bound CXCL17? Since constitutive expression of CXCL17 is restricted to gastric and respiratory mucosal tissues, CXCL17 expressed at such locations is perfectly placed to bind with strong affinity to GAGs present either in free forms or bound to cells. Such a process may permit the retention and oligomerization of CXCL17 on mucosal surfaces to create high local concentrations at pathogen infection routes. Given reports of bactericidal and fungicidal activity against a variety of pathogens [16], GAG-bound CXCL17 is likely to form a first line of defence against microbes. Consistent with such a function, CXCL17 expression by epithelial cells has been reported to be induced following challenge of mice with Mycobacteria, although CXCL17-deficient mice were found to be no more susceptible to infection than wild-type littermates [71]. CXCL17-deficient mice have also been reported to be less resistant to infection in a mouse model of herpes simplex virus infection, although this was postulated to be a result of impaired trafficking of GPR35^+^ cytotoxic T cells [72].

Mechanistically, cationic antimicrobial proteins exert their effects by associating to negatively charged lipopolysaccharides (LPS) on gram-negative bacterial membranes, where they introduce transient membrane defects or form stable pore-complexes to permeate the membrane, killing the pathogen (reviewed in [18; 73; 74]). Burkhardt *et al.,* previously reported CXCL17 exerts its antimicrobial effects on E.*coli* by permeabilising the bacterial membrane, and rightly speculated with the available structural models for CXCL17 at the time, that due to the constraints of the general chemokine structure the antimicrobial mechanism of action must differ from that of other linear antimicrobial peptides and defensins [16; 75; 76]. Here we provide evidence that the structure of CXCL17 is unlikely to be constrained in this manner and therefore raise the possibility that CXCL17 may partly function as a monomeric or dimeric pore-forming antimicrobial protein; a question which may now merit further investigation. Interestingly, the fungicidal activity of CXCL17 suggests it also has the capacity to permeabilise eukaryotic cell membranes, which could potentially be mediated by the charged carboxyl tail as similarly described for CCL28 [77].

Additionally, via the same electrostatic mechanisms by which CXCL17 is likely to associate to bacterial membranes, the cationic residues within CXCL17 may permit binding of LPS, thereby dampening innate immune responses originating in the mucosae. Moderately high concentrations of CXCL17 (300 nM) were shown to significantly reduce LPS-induced transcription of inflammatory markers IL-6, TNFα and iNOS in macrophages *in vitro*, in a manner that was enhanced by priming the cells in the presence of CXCL17 overnight [17]. While no mechanism was proposed for this inflammation suppressing effect at the time, cell surface GAG-bound CXCL17 could be envisaged to bind free LPS or remodel the macrophage glycocalyx to fine-tune TLR4 mediated responses.

Contrary to previous reports of chemotactic activity assigned to CXCL17, we could only demonstrate weak agonist activity for human neutrophils and failed to see convincing migratory responses of murine splenocytes. Previously, we and others have struggled to show robust chemotactic activity for monocytes and THP-1 cells using a commercially available CXCL17 (24-119) form [22; 23]. We therefore assumed that CXCL17 might play another physiological role. Indeed, at micromolar concentrations of CXCL17, we observed disruption of CXCL8-mediated chemotaxis of CXCR1 transfectants. We speculate that this may be due to competition for cell-surface GAGs on the migrating cell, or CXCL17 induced remodelling of the glycocalyx, since we have previously reported that GAG-binding is a requirement for chemotactic responses to CXCL8 [44]. A similar inhibitory observation was recently reported for the CXCL12-CXCR4 signalling axis [78], suggesting that CXCL17 may have broadly inhibitory effects on the function of a variety of chemokines and may serve to moderate chemokine signalling *in vivo.* A lack of such inhibition may explain in part, the perturbed trafficking of lymphoid and myeloid cells and exacerbated disease reported in CXCL17-deficient mice utilised in a model of experimental autoimmune encephalomyelitis (EAE) [79]. We can also speculate that while disrupting inappropriate pro-inflammatory signals originating in the gastric and respiratory mucosa, CXCL17 may still permit the transmigration of leukocytes via receptor independent GAG-mediated interactions as recently described for CXCL4 [56], potentially by condensing cell-associated heparan sulfate proteoglycans to generate a denser glycocalyx and promote cell-surface interactions with the local extracellular matrix; so while inappropriate anti-commensal immune responses may be dampened, tissue homoeostasis and immune sensing of the microbiome may still take place by patrolling leukocytes.

In summary, we report that CXCL17 displays some but not all the properties associated with a chemokine. CXCL17 has only feeble chemoattractant properties for human neutrophils, which may be due to the adoption of a protein fold not consistent with other members of the chemokine family. However, like some other chemokines, CXCL17 has a tendency to form dimers and is an efficacious binder to a variety of GAGs. GAG binding may enhance the anti-microbial of CXCL17 at mucosal surfaces, and it is possible that CXCL17 modulates the function of other chemokines dependent upon interactions with GAGs for their signalling. Peptides derived from the C-terminus of CXCL17 may therefore have immunomodulatory functions as has been exploited by others, for example C-terminal derivatives of CXCL9 [80; 81; 82; 83]. As such, CXCL17 may prove to be a useful tool with which to modulate a variety of inflammatory processes.

## 7 Author Contributions

JP conceived and designed the study. SG, SR, SH, HB and KM carried out the experimentation. TT and SK provided expert help with real-time chemotaxis assays. RS supported the work with mouse splenocytes. DD provided expert interpretation of BLI datasets. SG performed the statistical analyses and wrote the first draft of the manuscript. All authors contributed to manuscript revisions, read and approved the submitted version.

## 10 Data Availability Statement

Datasets are available on request. The raw data supporting the conclusions of this article will be made available by the authors, without undue reservation.

