## Supplementary materials for "CXCL17 binds efficaciously to glycosaminoglycans with the potential to modulate chemokine signalling"

### Supplementary Figures

**Supplementary Figure 1. Increasing the rate of AlphaFold2 structural model recycling maintains the prediction that CXCL17 (24-119) has a C-terminal alpha-helix.** The structure of CXCL17 was modelled with ColabFold: AlphaFold2 using various iterances of model recycling, as this has previously been described to improve the quality of structures for which low numbers of homologs are present in reference libraries. (**A**) Multiple sequence alignment (MSA) was performed against CXCL17 (24-119) with MMSeqs2 Uniref90 and environmental databases, with the sequence coverage per amino acid position of CXCL17 displayed. The structure of CXCL17 was modelled 3 times, either with 3 (**B-D**), 12 (**E-G**) or 48 (**H-J**) iterances of model recycling. Stereochemical plausibility and confidence in the models are reflected in the plDDT scores per residue position for the top 5 ranking models (**B, E, H**). Plots of predicted alignment error (PAE) per residue position for the top 5 ranking models are displayed (**C, F, I**). For each predictions with 3 and 12 iterations of model recycling, the rank 1 structure model is rotated 90˚ around the Y-axis (**D, G**). The top 5 highest ranking structural models of CXCL17 (24-119) generated with 48 iterances of model recycling are displayed (**J**). Secondary structure is indicated (α-helix; cyan, random coil; salmon pink), and all cysteines are coloured yellow.

**Supplementary Figure 2. Structural prediction of CXCL17 (24-119) by C-I-TASSER questions chemokine-like homology.** Structural predictions of CXCL17 (24-119) were generated by C-I-TASSER, integrating the I-TASSER hierarchical structure modelling approach with deep learning based contact predictions to guide Replica Exchange Monte Carlo (REMC) folding simulations. (**A**) Secondary structure predictions by amino acid position are presented where 'C' denotes random coil, and 'H' denotes α-helix. The top 5 ranking models by cluster size (Models 1-5); rank 1 (**B**) and ranks 2-5 (**C-F**), were rendered in pyMOL with secondary structure indicated (α-helix; cyan, random coil; salmon pink) and all cysteines coloured yellow. Confidence (C-scores) for each model are indicated. For **B**, all positively charged arginine and lysine residues were coloured blue, the model is rotated 180˚ and displayed without and with molecular surface. $NH_2$- and COOH- termini are annotated as N and C respectively.

**Supplementary Figure 3. Structural modelling of CXCL17 (24-119) with RoseTTAFold ColabFold using MMSeqs2 reference library or single sequence input for *de novo* folding**. (**A**) ColabFold RoseTTAfold structural model of CXCL17 (24-119) using MMSeqs2 and Uniref library (pdb70 template input). The MSA displays the sequence coverage of the input against homologs in the UniRef90 and environmental libraries. (**B**) Shows plDDT scores per residue, indicating local confidence in the model, and (**C**) shows the top ranking structural prediction generated in PyMOL. (**D**) ColabFold RoseTTAfold structural model of CXCL17 (24-119) using single sequence input settings for *de novo* folding of proteins with few homologs in the UniRef90 and environmental reference libraries. plDDT scores per residue are indicated, and (**E**) shows the top ranking structural prediction generated in PyMOL. All structures are shown with rotation 90˚ on the Y-axis, with all cysteines coloured yellow, all arginines and lysines coloured blue, and secondary structure elements coloured teal (α-helix) and salmon pink (random coil). $NH_2$- and COOH- termini are annotated as N and C respectively.

**Supplementary Figure 4. Structural modelling of CXCL17 (24-119) with AlphaFold ColabFold using single sequence input for de novo folding of proteins with low numbers of homologs in the reference library**. ColabFold AlphaFold2 structural model of CXCL17 (24-119) using the single input sequence setting for de novo folding of proteins with low numbers of homologs in the UniRef90 and Environmental reference libraries. No MSA was performed, (**A**) plDDT and (**B**) PAE scores per residue for the top 5 ranking models (rank1-5) are displayed. (**C**) The top ranking structural model was generated in PyMOL and rotated 90˚ on the Y-axis. All cysteines are coloured yellow, all arginines and lysines coloured blue, and secondary structure elements coloured teal (α-helix) and salmon pink (random coil). $NH_2$- and COOH- termini are annotated as N and C respectively.

**Supplementary Figure 5. AlphaFold2 ColabFold structural prediction of CXCL17 (24-119) homodimer displaying the top 5 ranking models.** Structural predictions of CXCL17 (24-119) homodimer were generated by AlphaFold2 ColabFold using 12 iterances of model recycling. (**A**) The sequence coverage from the initial MSA step is displayed and (**B**) stereochemical plausibility and confidence in the models are reflected in the plDDT scores per residue position for the top 5 ranking models. (**C**) Plots of predicted alignment error (PAE) per residue position for the top 5 ranking models are displayed. (**D-H**) Corresponding structural predictions for models ranked 1-5 respectively. Predicted structures are displayed with all cysteines coloured yellow and secondary structure elements coloured teal (α-helix), magenta (β-sheet) and salmon pink (random coil). $NH_2$- and COOH- termini are annotated as N and C respectively.

**Supplementary Figure 6. Production of recombinant In-house CXCL17 (24-119) .** (A) 6xHis-SUMO3-CXCL17 (24-119) was purified from the soluble fraction of Shuffle T7 Express *E.coli* by IMAC with the resulting HPLC chromatogram displaying $A_{280nm}$ absorbance and (B) SDS-PAGE with Coomassie stain displaying the distribution of total protein eluted in various fractions of the purification. The 5 right-most lanes of the SDS-PAGE gel image show the distribution of protein at various stages of the digestion procedure of 6xHis-SUMO3-CXCL17 (24-119) with *Ulp1*. (**C**)

Solubilized and refolded CXCL17 (24-119) was purified by HiTrap heparin HP 1ml affinity chromatography and (**D**) total protein eluted from the column was visualized by SDS-PAGE with Coomassie stain. Recombinant in-house generated proteins were resolved by Western blot with CXCL17 anti-serum (sheep anti-human pAb (AF4207). (**E**) serial dilutions of SUMO3-CXCL17 (24-119), (**F**) and CXCL17 (24-119) . Dimer complexes were covalently crosslinked by addition of 5mM $BS^3$ chemical crosslinker to various concentrations of (G) SUMO3-CXCL17 (24-119), (H) and CXCL17 (24-119). Images are representative of n=3 separate experiments. Protein identity was also confirmed by probing with mouse anti-hCXCL17 mAb (MAB4207) (data not shown).

**Supplementary Figure 7. The SUMO3 tag does not interfere with CXCL17 binding to Heparin, HS or CS.** The N-terminal SUMO3-tag on SUMO3-CXCL17 (24-119) and the C-terminally truncated mutants was confirmed not to interact with binding of CXCL17 (24-119) to GAGs using a colorimetric static-binding assay. Preparations of CXCL17 (24-119) , SUMO-CXCL17 (24-119) and SUMO3-tag were immobilised, to which the binding of 1µg/ml biotinylated GAGs heparin (**A**), HS (**B**) or CS (**C**) were quantified (n=4). Data were normalized to maximum binding recorded per experiment and display mean ±SEM. (**D-F**) 500nM CXCL17 (24-119) and SUMO-CXCL17 (24-119) were immobilised in both their native state and after being reduced with 2mM DTT, to which the binding of 1µg/ml heparin, HS or CS was quantified (n=3). Data displayed as mean +SEM. Two-way ANOVA with multiple comparisons and Dunnett's post-test was performed and no statistical significance was detected (NS).

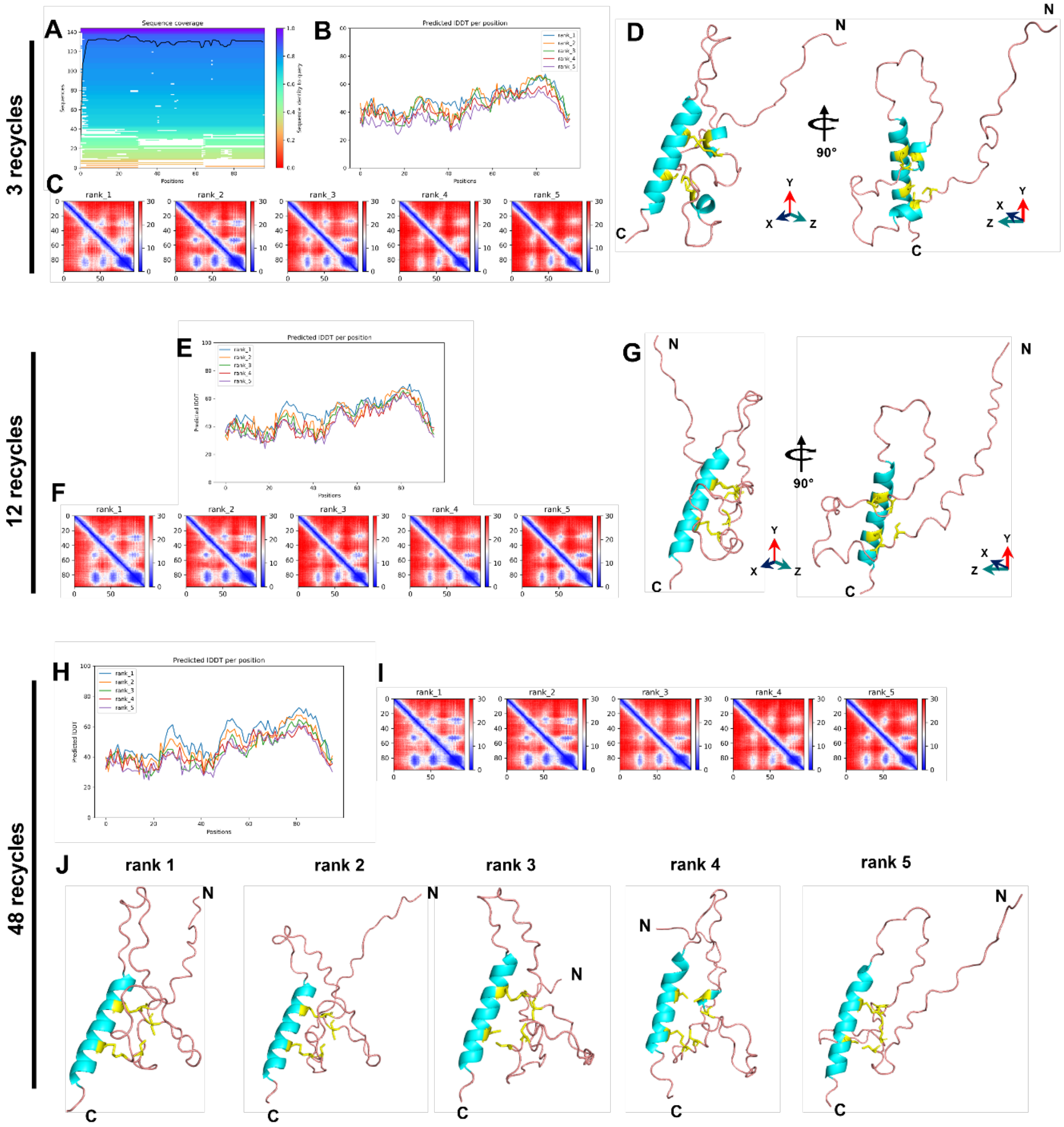


**Supplementary Figure 1.**

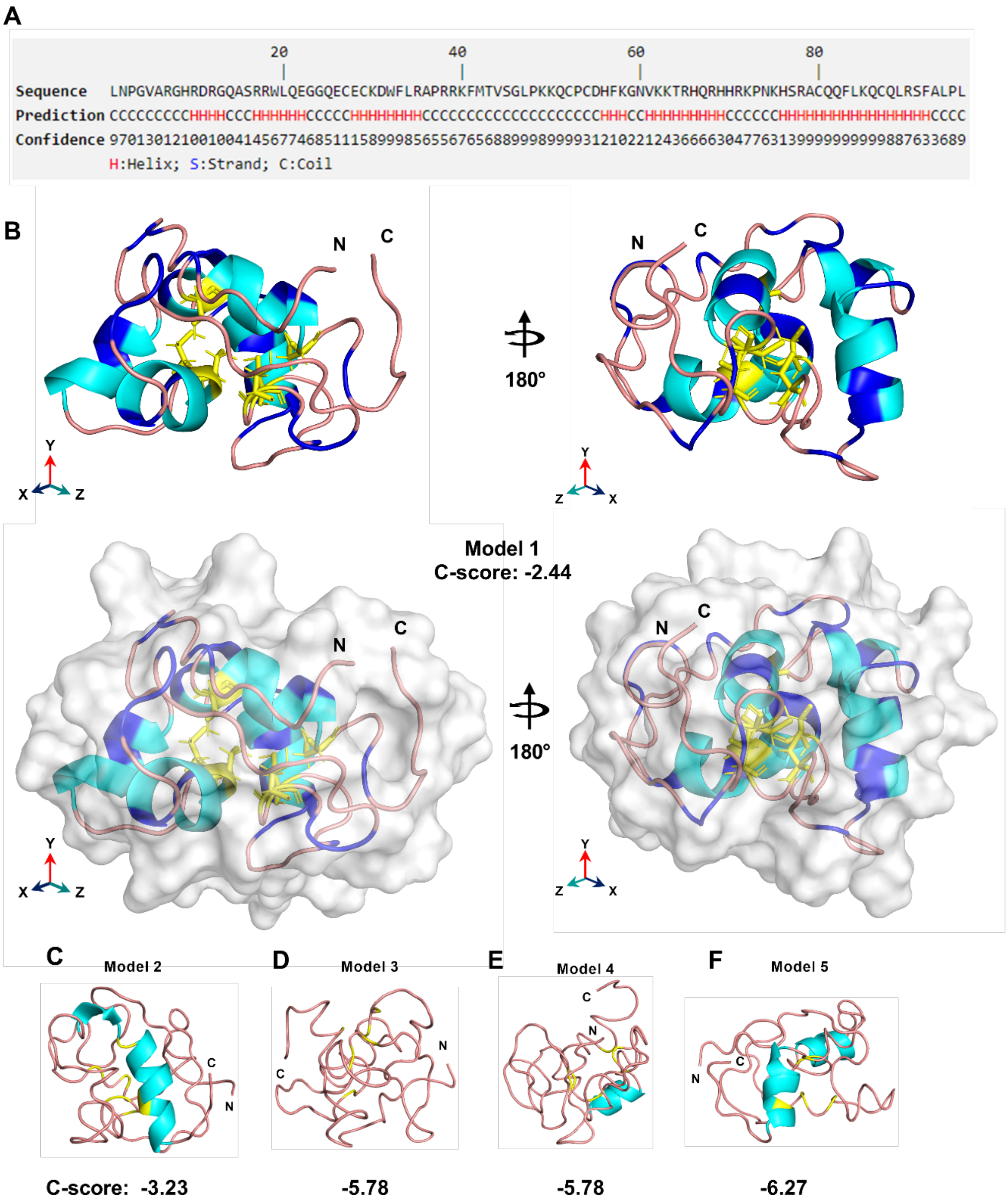


**Supplementary Figure 2.**

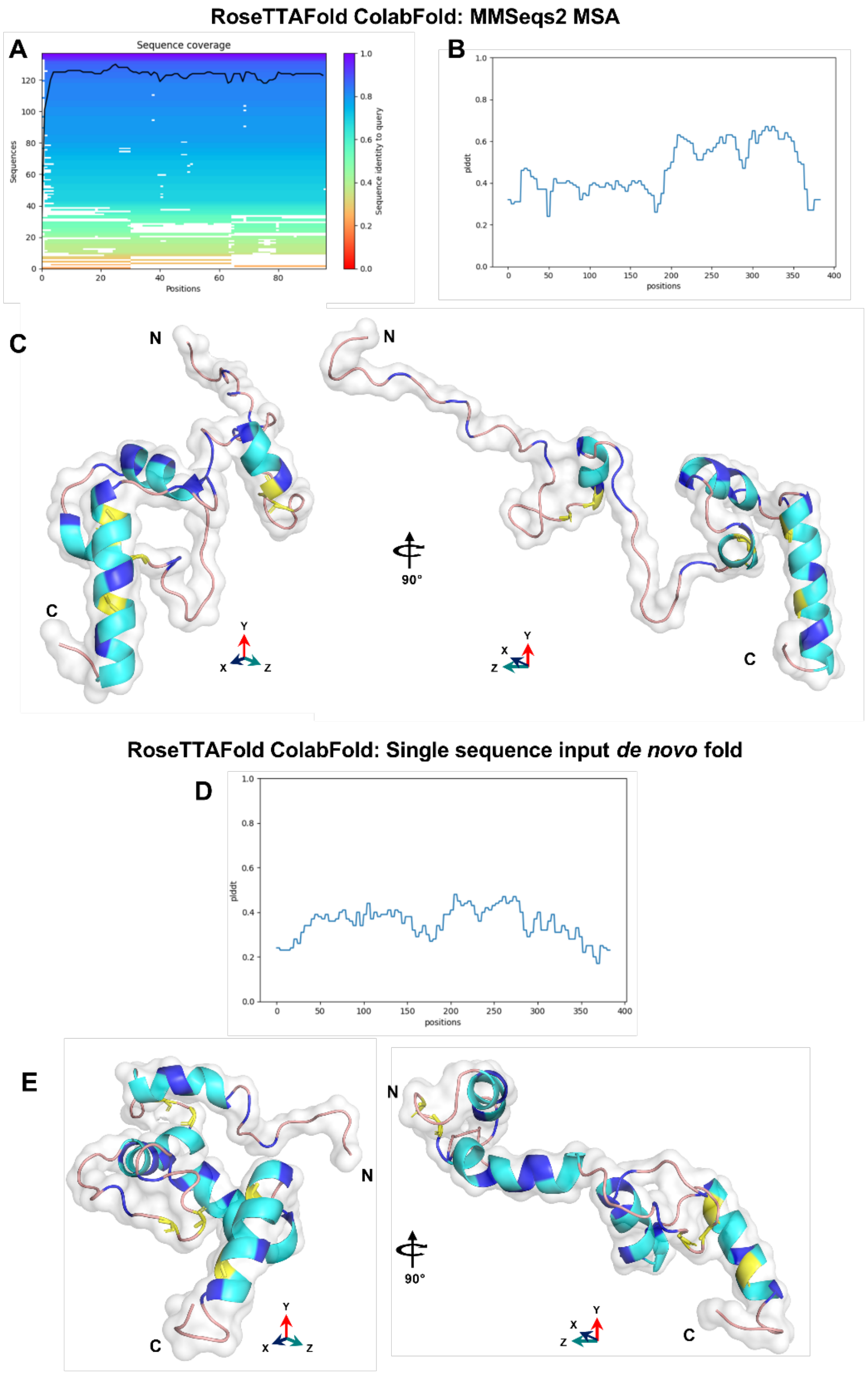


**Supplementary Figure 3.**

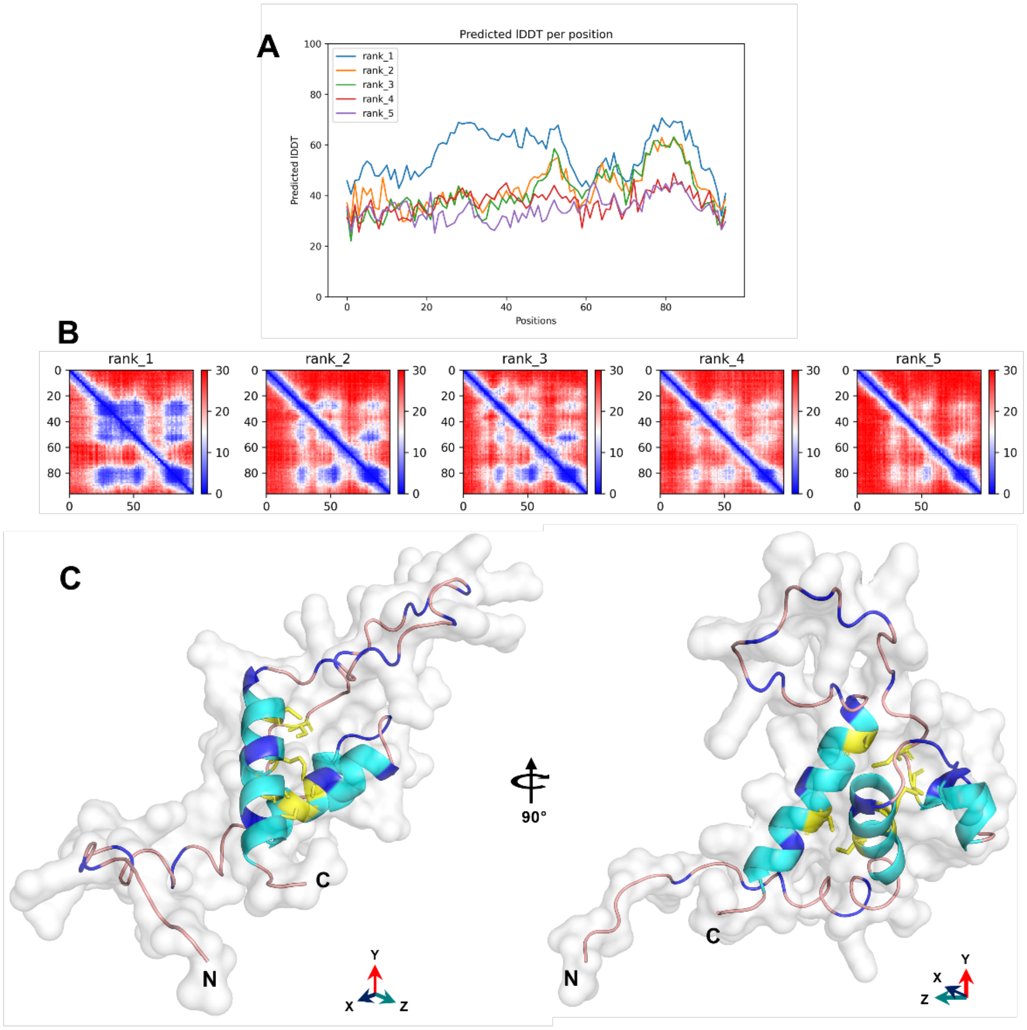


**Supplementary Figure 4.**

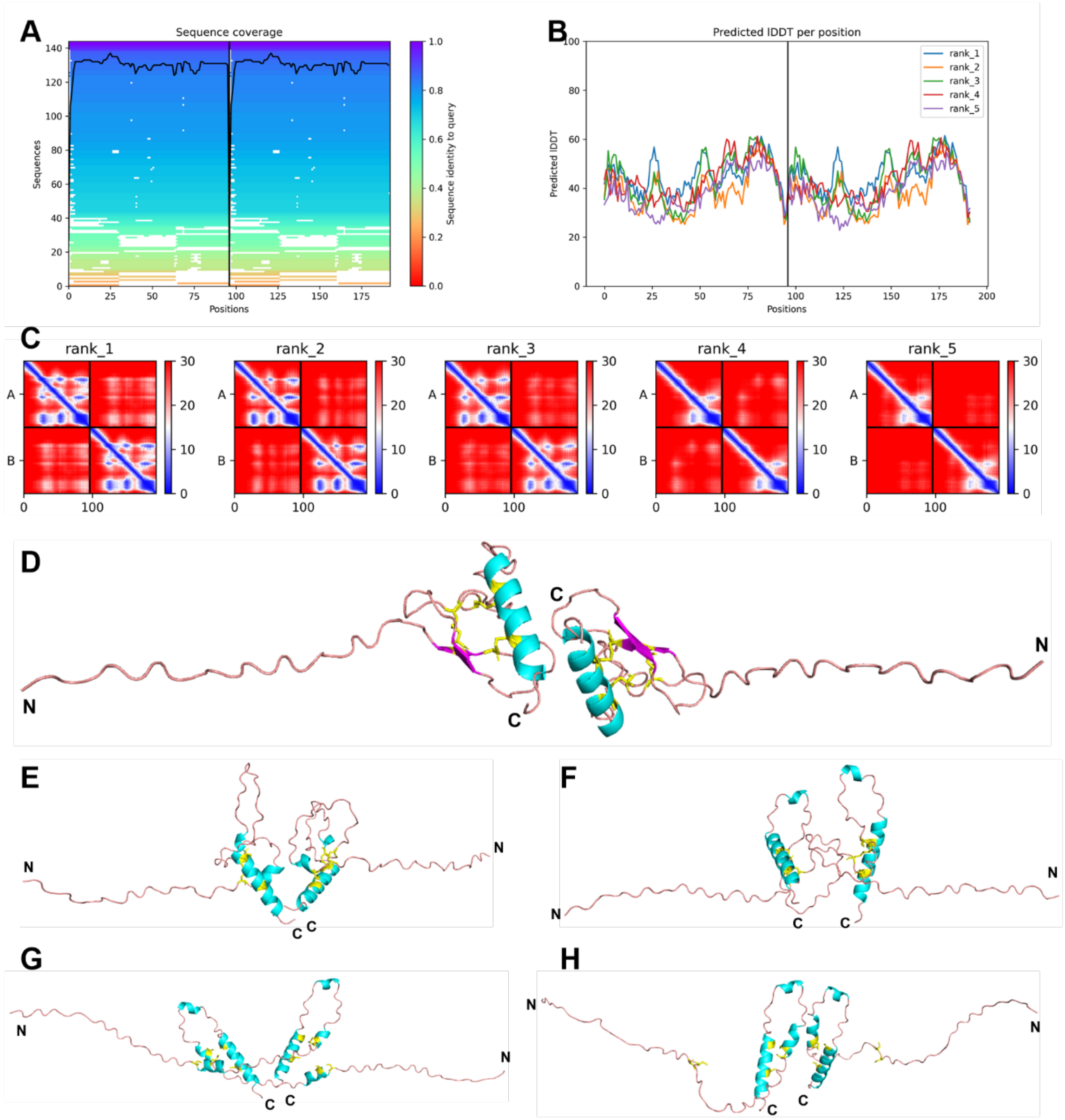


**Supplementary Figure 5.**

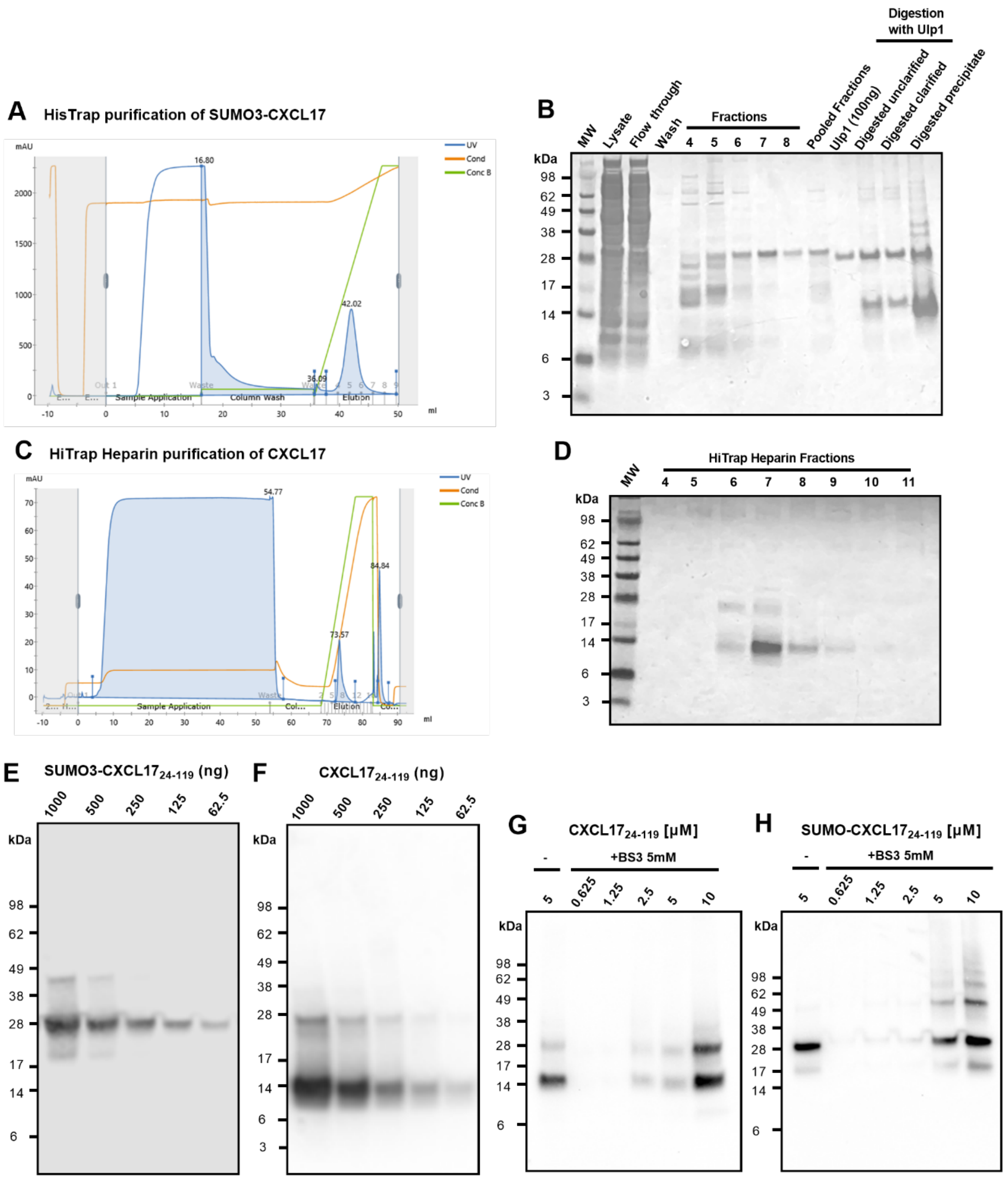


**Supplementary Figure 6.**

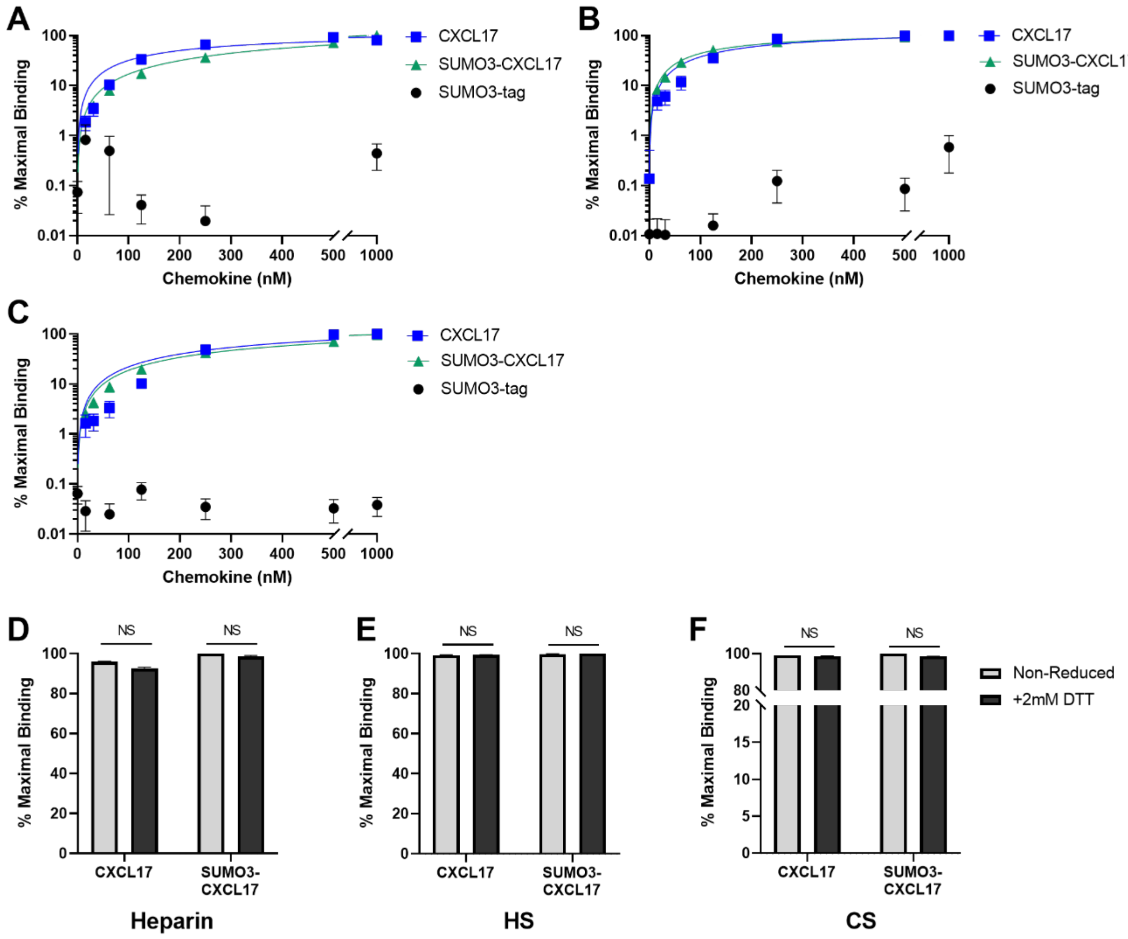


**Supplementary Figure 7.**
